# The LOTUS-domain containing protein PpLDCP3 controls germline and dispersal unit formation in the moss *Physcomitrium patens*

**DOI:** 10.1101/2024.09.17.613224

**Authors:** Anton Kermanov, Jörg D. Becker

## Abstract

- Dispersal is crucial for the survival and thriving of plant populations, yet the mechanisms of dispersal unit formation are poorly understood. LOTUS domain-containing proteins are essential for animal reproduction, promoting cell cycle control, transposon silencing, and transgenerational inheritance in the germline. In this study, we demonstrate that the formation of dispersal units, sperm and spores, in the model moss *Physcomitrium patens* relies on the novel LOTUS domain-containing protein 3 (PpLDCP3).
- *Ppldcp3* knock-out mutants revealed pleiotropic phenotypes in both gametogenesis and sporogenesis. *Ppldcp3* sperm were rarely released and appeared to be partially non-viable, leading to fewer sporophytes produced. Sporophyte development in *Ppldcp3* was impaired, resulting in abnormal dehiscence of spores, which were incapable of germinating.
- Fluorescent reporter lines revealed the presence of PpLDCP3 in developing gametangia. Within the premature egg cell and ventral canal cell of archegonia, PpLDCP3 was identified as a component of cytoplasmic fluorescent foci, which we interpret as P-bodies. This suggests a role for PpLDCP3 in RNA metabolism, analogous to animal LDCPs. Moreover, bioinformatic predictions indicate PpLDCP3 as a downstream player of WOX13 and bHLH transcription factors.
- We conclude that PpLDCP3 is integrated into evolutionary conserved mechanisms of dispersal unit formation in land plants.

## Introduction

The success of a species in ecological and evolutionary terms is essentially defined by its ability to spread genes. The sessile vegetative lifestyle of land plants brings the perils of genetic diversity decline due to inbreeding and increases the risks of habitat fragmentation (Baker, 1959; Charlesworth, 2003). The production of dispersal units, or propagules, can be considered a way to overcome this limitation (Robledo-Arnuncio *et al*., 2014; Saastamoinen *et al*., 2018; Gelmi-Candusso *et al*., 2019).

In seed plants, the dispersal is facilitated by two key mechanisms: pollen grains and seeds (Pacini, 2012). Dispersal of pollen grains potentiates fertilization of immobilized egg cells located on a remote or the same plant shoot by pollen tube-delivered sperm cells (Hafidh & Honys, 2021). Seeds give rise to plants with distinct genetic backgrounds at a new reproduction site, promoting the gene flow in a population (Robledo-Arnuncio *et al*., 2014).

However, during land plant evolution, cryptogams were the first to face terrestrial environments and had to evolve strategies for dispersal that differed from their Streptophyte ancestors (Niklas & Kutschera, 2010; Jill Harrison, 2017). The innovations can be exemplified by two dispersal stages incorporated into the life cycle of bryophytes. In the extant representative *Physcomitrium patens*, dispersal events terminate the development of two distinctive generations of the multicellular gametophyte and sporophyte. On a *P. patens* gametophyte, sperm matures in male reproductive organs (antheridia), surrounded by sterile jacket cells, and is released into the environment through an opening in the antheridial tip. Sperm is then actively moving in surface water, which is a prerequisite for the fertilization of an egg cell, embedded into a multilayered egg cell chamber of an archegonium, located on the same or different shoot (Cove, 2005).

A single, non-branching diploid sporophyte develops from a fertilization event, resulting in the production of desiccation-tolerant haploid spores covered by sporopollenin. These spores are released through a crack in the spore capsule, marking the end of the sporophytic generation (Cove, 2005; Rensing *et al*., 2020). The dispersal of dry sporopollenin-covered spores confers resistance to various terrestrial abiotic factors, including prolonged UV irradiation and dehydration, and protection from terrestrial predators (Delwiche *et al*., 1989; Becker & Marin, 2009).

A few thousand sperm per antheridial bundle and a comparable number of spores per fertilized shoot ensure proper dispersal in both water and dry environments (Engel, 1968; Rensing *et al*., 2020). Formation of these dispersal units must be ensured by their synchronized development, controlled maturation, and release as separated units from the plant body. The formation of a multicellular antheridium and sporophyte, “nursing” these cells, represents an evolutionary novelty in bryophytes and required both the appearance of new genes and the rewiring of pre-existing gene networks.

In animals, several proteins with a domain called LOTUS (***<u>L</u>****imkain*, ***<u>O</u>****skar*, and ***<u>Tu</u>****dor* containing proteins 5 and 7) (Callebaut & Mornon, 2010) or OST-HTH (***<u>Os</u>***kar-***<u>T</u>***DRD5/TDRD7 ***<u>HTH</u>***) (Anantharaman *et al*., 2010) domain were shown to be essential for reproduction. The unifying feature of these proteins is the expression in the germ cells and surrounding nursing cells, regulation of the cell cycle by affecting cyclin abundance, reproductive genome safeguarding from transposon activity, and selective clearance of mRNA transcripts through interaction with deadenylation and decapping complexes (Kubíková *et al*., 2021). In animals, the appearance of specific LOTUS domain architectures correlates well with the radiation events, leading to the appearance of phylogenetically new clades, such as in the case of Oskar in Holometabola (Blondel *et al*., 2021) and TDRD5/7 in metazoans.

Here we describe the LOTUS-domain containing protein 3 (LDCP3) with a novel domain architecture, appearing in early land plants. We establish that PpLDCP3 bears functions in the late stages of gametogenesis and sporogenesis of *P. patens*, providing dispersal capacity for both gametophytic and sporophytic generations. We reveal parallelisms between the PpLDCP3 and animal LDCPs functions from a developmental perspective.

## Materials and methods

### Plant growth conditions

The wild-type Gransden line of *Physcomitrium patens* Bruch & Schimp (Ashton & Cove, 1977) was used in this study.

### Plant growth on media

Plants were grown on sterilized KNOPS medium (Reski & Abel, 1985), with the addition of 0.5 g/l ammonium tartrate dibasic and 5 g/l glucose, and solidified by the addition of agar to 1% final concentration. Protonema was subjected to grinding every 1-2 weeks with tissue rupture in sterile water and placed on a sterile fresh medium, overlayed with sterile cellophane. Spore germination assays were carried out on the same medium with cellophane overlayed. For transformant selection, the medium was supplemented with antibiotics in appropriate concentrations: hygromycin B (30 µg/ml) for the selection of *Ppldcp3* and G418 (25 µg/ml) for the selection of PpLDCP3 translational fusion lines.

### Plant growth on soil

Autoclaved rehydrated peat pellets (Jiffy-7; Jiffy Products) (referred to in the text as “jiffies”) were used to grow plants from protonemata or spores in corresponding “long day” conditions, and reproductive organ formation was induced by short-day conditions as described elsewhere (Ortiz-Ramírez *et al*., 2017). For more details, see supplementary methods.

### Generation of *P. patens* mutant lines

#### Ppldcp3 full knock-out mutant construct generation

pBHRf vector was modified to carry the right and left flanks of the Ppldcp3 gene, with a Hygromycin resistance cassette in between, as described elsewhere (Schaefer *et al*., 2010).

#### PpLDCP3 translational fusion constructs generation

Translational fusion lines were created using Gateway technology with a system with four-fragment recombination vectors and pGEM-gate as destination vectors, all kindly provided by M. Bezanilla’s lab. All translational fusions are C-terminal to the PpLDCP3 protein sequence.

Both deletion and translational fusion lines were obtained by homologous recombination, occurring after the transformation of wild-type protoplasts with corresponding linearized plasmids. For more details, see supplementary methods.

### Phenotyping assays

#### Mature sporophyte counts

Mature sporophytes were counted at 60-70 days post-induction on all the shoots. At least 200 shoots were used for each counting for wild-type and *Ppldcp3* knock-out mutant plants. At least 25 shoots were observed for the PpLDCP3 translational fusion lines. An average of 3-5 measurements with standard error is presented in the corresponding table.

#### Sporophytes opening measurement

Sporophytes on the 90-100 dpi shoots were collected and examined under a stereoscope. Opened sporophytes were detected as the ones containing seta or leftover sporophytic tissue/spores (group “opened”, “O”). Unopened sporophytes were gently touched using tweezers, sufficient for spore dehiscence in some cases (group “opened by touch”, “T”). Sporophytes that could release spores only by prolonged pressure applied by tweezers or squashing were referred to as “opened by applying pressure”, or “P”. At least 20 shoots with sporophytes were observed for a single measurement and the percentage of sporophytes belonging to each group was calculated. The averages of multiple measurements for each group were plotted as bar plots.

#### Imaging of crushed live sporophytes

Live sporophytes were collected from 60-70 dpi shoots and crushed by pressing the coverslip against the sporophyte on a microscopy slide on the same day and used for imaging.

#### Spore germination assay

Mature sporophytes without visible cracks were collected in batches, with each batch consisting of 3 spore capsules. Sporophytes were stored in the fridge (+4 °C) for no longer than 2 months or processed immediately. Sporophytes were incubated for three minutes in bleach (diluted with water 1:2), then washed with sterile water, and opened with a sterile filter tip in 300 ul of water. Of this volume, 200 ul was aliquoted and plated on soil; while 100 ul was supplemented with 900 ul of water, and 300 ul was plated on plates with KNOPS-T solid media, 3 replicates for each genotype. Germination of spores was observed visually every 5 days until 20 days post-plating. Germination on soil was observed for 50 days.

#### Spore counting

Mature sporophytes were fixed in FAA solution (formaldehyde: acetic acid: alcohol (1:1:18)). Spores were released in water and counted using a hemacytometer. For more details, see supplementary methods.

#### Histochemical sectioning and staining

Sporophytes for histological sectioning were collected from wild-type and *Ppldcp3* plant plants at the late developmental stages and fixed in FAA (formaldehyde: acetic acid: alcohol (1:1:18)) solution. After sample processing, slides were stained for 5 minutes in 1% TBO with 2% Sodium Borate, washed with PBS+T, and imaged. For details of the protocol, see supplementary methods.

#### Live-dead staining assay

For each observation, 10 shoots were dissected from the wild type and *Ppldcp3*. Gametangia were then placed in 20 μl of sterile water on a microscopy slide, divided into 2 sections (wild type/mutant), and covered with a coverslip. The slide was placed under the microscope and 2 ul of both SYTO9 and propidium iodide from LIVE/DEAD™ BacLight™ Bacterial Viability Kit (previously diluted 1:20 *ex tempore* in water) were slowly added from the edge of the coverslip and mixed by pipetting (total dilution approximately 1:200). Slides were placed on an orbital shaker for 10 minutes covered by aluminium foil and then observed under the microscope.

### Imaging

#### Photographing

Protonema on KNOPS-T plates, germination of spores, jiffies with developed plants, and individual gametophores were imaged using a Nikon D7500 camera. For wild-type and *Ppldcp3* gametophores imaging, 3 representative gametophores of each genotype, grown in standard conditions on jiffies, were taken at 50-60 days dpi. For sporophyte imaging and subsequent size measurement, at least 3 different jiffies per genotype were used for the sporophyte collection and fixation in batches. Sporophytes were imaged using a DFK 24UP031 color camera controlled by IC Capture 2.5 software.

#### Histochemical imaging

Slides with the stained histochemical sections of sporophytes were imaged with a Hamamatsu Digital scanner C13140-01.

#### Bright-field and epifluorescence microscopy

Imaging of spores, crushed sporophytes, and sperm observations were performed using a Leica DM6 B, equipped with a color camera and controlled by LasX software. Epifluorescence imaging of live-dead staining assay, as well as of LDCP3-mCherry/mEGFP gametangia, was performed on the same device with excitation/emission settings for mEGFP and mCherry of 480/527 nm and 546/605 nm, respectively. Microscope settings were adjusted to reduce plant tissue autofluorescence. Imaging of fluorescent sporophytes was performed with a stereoscope Zeiss Lumar with a HamamatsuHam DCAM camera.

### Confocal microscopy

Confocal Z-series stacks were acquired on a Zeiss LSM 880 point scanning confocal microscope using photomultiplier tube detectors (PMTs) and a gallium arsenide phosphide (GaAsP) detector, 10x EC Plan-NEOFLUAR and x20 Plan-Apochromat 1.4NA (0.3/0.8 NA correspondingly) air objective (Zeiss) and the 488 nm and 561 nm laser lines. Zeiss Zen 2.3 (black edition) software was used to control the microscope and adjust spectral detection.

### Video recording

A video recording of live *Ppldcp3* sporophyte dissection was performed using a stereoscope Nikon SMZ800, and an attached digital camera with a 12MP sensor with 1.22 µm pixels and a 28mm f/1.8 stabilized lens. Sperm movement, observed using a Leica DM6 B, was recorded using the ezvid 1.0.0.4 program.

### Image processing and analysis

Histochemical section images were processed using the NDP.view2 program with adjustments to scale. All other images and videos were analyzed with FIJI (ImageJ2), version 2.9.0/1.53t. Fluorescent Images of wild-type and translational fusion lines (“paired” images) on the corresponding panels were adjusted for brightness and contrast with equivalent settings.

### Bioinformatics

For the prediction of the PpLDCP3 interaction network, protein structure modeling and BLAST searches for the longest isoform of PpLDCP3 were used. For SmartBlast searches (https://blast.ncbi.nlm.nih.gov/smartblast) prefiltering of the output was done to limit it to non-plant representatives, with alignment of the selected candidates using COBALT (https://www.ncbi.nlm.nih.gov/tools/cobalt/re_cobalt.cgi). AT2G05160 predictions were taken from automatic ThaleMine predictions (https://bar.utoronto.ca/thalemine/report.do?id=4537526&trail=%7c4537526).

### Data handling, statistical analysis, and graphics

Data analysis was performed using in-house developed Python scripts in IDE PyCharm 2023. Indication of statistical significance with stars corresponds to values:

ns: 5.00e-02 < p <= 1.00e+00

*: 1.00e-02 < p <= 5.00e-02

**: 1.00e-03 < p <= 1.00e-02,

***: 1.00e-04 < p <= 1.00e-03

****: p <= 1.00e-04; where p is the P-value.

For box plots, the center line is the median; box limits are upper and lower quartiles; whiskers are minimum and maximum values. For point plots and strip plots, a horizontal black marker line corresponds to the average value. For line plots, vertical bars represent the standard error.

For all types of graphs, points are individual values, obtained as the result of independent measurement, superimposed on graphs.

Pictograms depicting *P. patens’* developmental stages were developed using BioRender. Genetic maps were generated using SnapGene v 3.1.4.

## Results

### *Ppldcp3* mutant sporophytes develop abnormally and produce non-germinating spores

In plants, the LOTUS domain is found in different protein architectures specific to their clades, distinct from those observed in animals (Anantharaman, Zhang, and Aravind 2010). Bioinformatic screening together with RNA-Seq data for *P. patens* (Julca *et al*., 2021) allowed us to identify a few **ldcp** (LOTUS domain-containing protein) genes of interest, expressed in the premature gametangia and sporophytes. One of these genes, named Ppldcp3, is expressed ubiquitously at low basal levels in *P. patens*, with peaks of expression corresponding to reproductive tissue in the gametophyte (i.e. premature antheridia and archegonia), and in premature sporophytes (green, meiotic-postmeiotic) (Fig. 1a). These stages preclude the formation of the moss dispersal units, represented by antherozoids and spores, respectively. Noteworthy, despite the expression during the formation of terminal cells, Ppldcp3 expression *per se* was not detected at significant levels in the sperm cells or the spores.

**Fig. 1.**
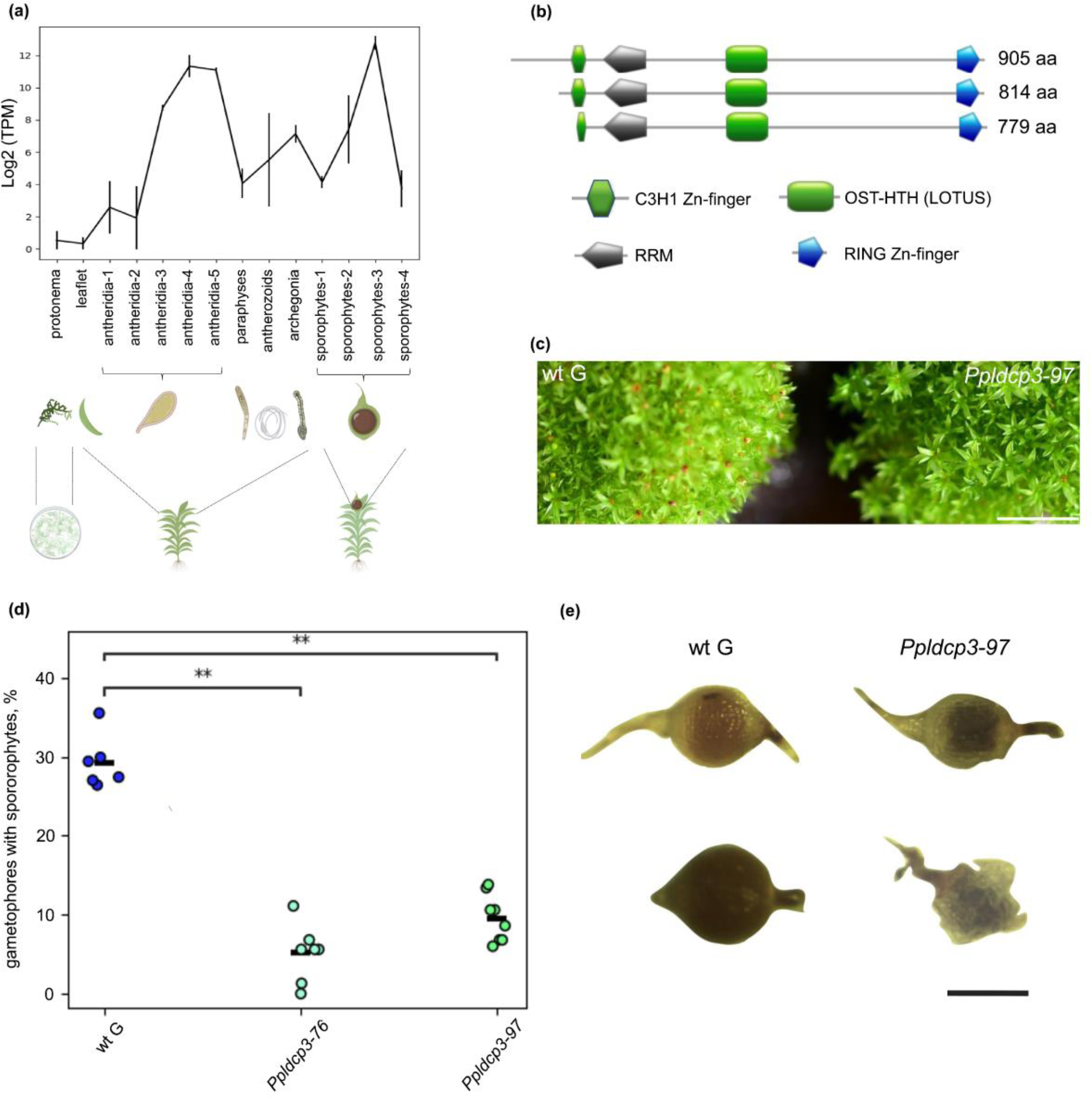
Ppldcp3 is expressed during the terminal stages of gametophyte and sporophyte development, and knock-out plants produce significantly lower amounts of sporophytes (a) Log2 transformed data of Ppldcp3 expression based on RNA-Seq data from the EVOREPRO database (https://evorepro.sbs.ntu.edu.sg/). Icons represent the late developmental stage of each category. The icons are not to scale. (b) Domains identified by ScanProsite tool (https://prosite.expasy.org/) for different polypeptide chains translated from Ppldcp3 locus in agreement with predicted transcript variants. Domain names, regions, and accessions as referred to in PROSITE, in the direction from N- to C-termini: C3H1 Zinc-finger type domain (PS50103), Eukaryotic RNA recognition motif (RRM) (PS50102), OST-HTH (PS51644), RING-type Zinc-finger domain (PS50089). OST-HTH corresponds to the LOTUS domain (cd08824) in the Conserved Domain Database (https://www.ncbi.nlm.nih.gov/Structure/cdd/cddsrv.cgi?uid=cd08824). Note that the shortest protein isoform (lower panel) has a reduced C3H1 Zinc-finger domain. (c) Image of *P. patens* gametophores grown on soil (jiffies), 70 days after transferring to short-day conditions. The sporophytes are visible on top of the gametophores. The *Ppldcp3-97* mutant line produces visibly fewer sporophytes than wild-type plants. The scale bar is 1 cm. (d) *Ppldcp3* mutant lines demonstrate a significant reduction in the number of sporophytes at 50-60 days post-induction. Each dot corresponds to the independent counting of sporophytes developed on 200 shoots. Statistical inference was based on the Mann-Whitney-Wilcoxon test, two-sided with Bonferroni correction. (e) *Ppldcp3* mature sporophytes (right panel) are polymorphic with abnormalities in development compared to wild-type ones (left panel). Distinct types of sporophytes in *Ppldcp3* were observed, some abortive in premature (mature-like) stages (top mutant spore capsule) often shaped irregularly, while others with amorphic protrusions of tissue. The scale bar is 500 µm.

Ppldcp3 encodes at least three protein isoforms, with the longest predicted to be 905 amino acids long. The domain architecture of the protein includes the Zinc-finger C3H1-type, RNA recognition motif (RRM), LOTUS (OST-HTH), and RING-finger domains (Fig. 1b). To get insights into Ppldcp3 function, we performed gene knock-out by homologous recombination (Fig. S1a). Five mutant lines passed all levels of selection on an antibiotic-containing medium. Two of those lines from independent transformations, which were confirmed to contain the insertions by PCR (Fig. S1b) and sequencing of the locus, were used for further studies.

*Ppldcp3* protonema (on KNOPS-T medium) and gametophores, grown both on medium and jiffies, developed normally without delays and visible differences in vegetative tissue morphology (Fig. S1c). Placing plants grown on jiffies into conditions triggering the development of reproductive tissue, however, resulted in a significantly lower number of mature sporophytes 60-70 days post-induction (dpi) in comparison to wild-type (Figs 1c, d). The low number of sporophytes produced in mutant plants might point to their parthenogenetic origin and hence a complete absence of fertilization events (Tanahashi *et al*., 2005). We tested whether supplementation of water on top of 15-19 dpi gametophores bearing mature antheridia would increase fertilization rates. We observed increased production of sporophytes on both wild-type and mutant plants, supposedly due to the gained capacity of sperm to be released from antheridia and fertilize egg cell, thus excluding the parthenogenetic explanation (Fig. S1d).

The *Ppldcp3* sporophytes developed normally until the meiotic stages. When a round sporophyte was formed, the difference between wild-type and mutant organs started to accumulate and became significant at the terminal stages (Fig. S1e). The majority of mature sporophytes, collected around 2-2.5 months post-induction were of abnormal shapes. Among the developmental effects observed in mutants were mixed-staged sporophytes, with the shape corresponding to the earlier developmental stages, but the color of capsules and visible spore-like content characteristic of the terminal stages in wild-type plants. Additionally, shrunken sporophytes, “budding”-like sporophytes, and sporophytes with green or yellow tissue protrusions derived from spore capsules were observed (some of these defects are shown in Fig. 1e).

The spore content of both wild-type and Ppldcp3-97 mutant mature sporophytes was also different. In preparations of smashed mature sporophytes, wild-type spores looked normally developed and properly shaped (Fig. 2a, upper panel). Quite the opposite, both wild-type-like and abnormally shaped mutant sporophytes appeared to consist of two populations of cells. One was represented by spore-like-looking cells, although smaller in size (Fig. S2) and number (Fig. 2d) than wild-type spores, while another consisted of dense particles, probably aborted cells, varying in size and shape. The latter was inconsistent and represented as non-globular, flattened, angular geometric forms (Fig. 2a, lower panel).

**Fig. 2.**
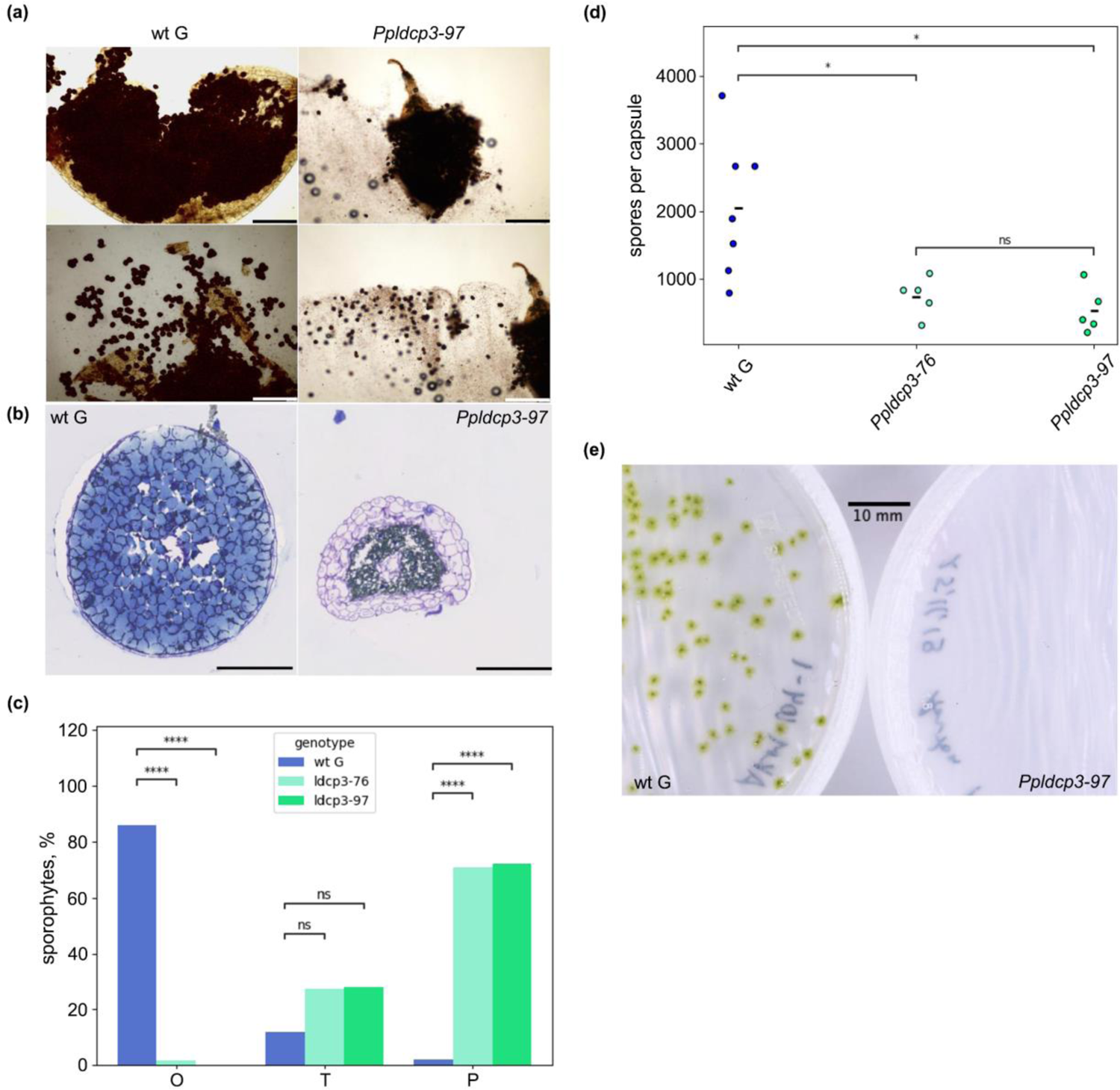
*Ppldcp3* sporophytes have developmental defects and do not release viable spores. (a) Wild-type and mutant sporophytes differ in their content. The squashing of sporophytes on a microscopy slide released dry mature spores in wild-type. In *Ppldcp3-97*, a heterogeneous cell population was released, together with the liquid it contains, comprising bigger spore-like cells and smaller particles. Elongated and “swollen” calyptra were visible on mutant spore capsules. Corresponding upper and lower panels represent the same spore capsules, but smashed further to stimulate spore spreading on the surface. The scale bar is 200 µm. (b) Histochemical staining with toluidine blue of transversal sections of wild-type and mutant mature sporophytes showed the presence of highly vacuolised amphithecium cells in *Ppldcp3-97* sporophytes, along with an irregularly shaped spore mass present inside the sporophyte. The scale bar is 125 µm. (c) Quantification of differences in dispersal unit release in a sporophytic generation. 90-100 days after induction most of the mutant sporophytes were still found unopened, while almost all the wild-type sporophytes showed cracked capsules or just leftovers of capsules, with dry spores released (“O”, opened) or being easily released by touch (“T”, touch). Most of the *Ppldcp3* sporophytes required squashing to release the spores (“P”, pressed). The averages and standard errors were calculated from at least 5 independent observations, with each observation including a minimum of 20 sporophytes observed. Statistical testing was performed using t-test for independent samples with Bonferroni correction. (d) Number of spores per mature sporophyte for wild-type and *Ppldcp3* mutant lines. (e) Spores collected from *Ppldcp3-97* mutant plants placed on KNOPS-T media were not germinating, unlike normally germinating wild-type plant spores. The scale bar is 1 cm.

These observations were supported by transversal histological sectioning and histochemical staining assays of fixed wild-type and *Ppldcp3-97* late-stage sporophytes, selected based on the absence of visual signs of abnormal sporophyte development (Fig. 2b). The pronounced layers of the amphithecium, which is a typical feature of earlier developmental stages in wild-type sporophytes, were consistently observed in the mature *Ppldcp3-97* mutant sporophyte. This was in contrast to wild-type mature sporophytes from the corresponding stage, in which the amphithecium represented a thin dry cell layer, ready to rupture and release mature spores. Amphithecium cells of *Ppldcp3-97* plants were full of vacuoles, suggesting perturbation of sporophyte desiccation and maturation, despite the overall mature-like morphology and dark-brown color.

Prolonged observations revealed defects in mutant spore dehiscence. At around 3 to 3.5 months post-induction, when mature wild-type sporophytes are mostly found opened and releasing mature dry spores, *Ppldcp3* sporophytes were stalled in their development, with no signs of the opening of the capsule. Only pressure applied to the sporophyte could trigger the spore release, producing the abovementioned mixture of spore-like particles together with accompanying aborted cells, all still embedded in a viscous liquid content (Video S1, Fig. 2c). We found that the ability to release the spores was to a large extent restored when sporophytes were grown submerged in water since the moment of fertilization (data not shown). This might imply a complex phenotype, related to the balance between water availability/conductivity and preservation of osmotic pressure in the developing sporophyte.

Spores from both wild-type and mutant sporophytes were tested for germination on KNOPS-T media (Fig. 2e). None of the spores of *Ppldcp3-97* plants germinated in 4 independent tests, indicating a complete lethality of the *Ppldcp3* genotype in the sporophytic generation.

### Ppldcp3 affects sperm quality and sperm release from antheridia

Previously described observations were related to sporophytic developmental defects and explained the observed phenotype to a rather limited extent. However, our previous observations with supplementation of water at the time of sperm release indirectly indicated a male reproductive phenotype (see Fig. S1d).

Therefore, we decided to observe the release and quality of sperm in *Ppldcp3-97*. In live-dead staining, antherozoids released from wild-type antheridia were mostly found to be viable (green fluorescent channel), while in the *Ppldcp3-97* mutant line a large proportion of sperm cells was found as dead cells in the red fluorescent channel (Fig. 3a). It is worth noting that the population of mutant antherozoids appeared to be partially dead even before release from the sperm matrix or soon afterward, and the number of dead cells gradually increased over time even before the cells were able to leave the matrix.

**Fig. 3.**
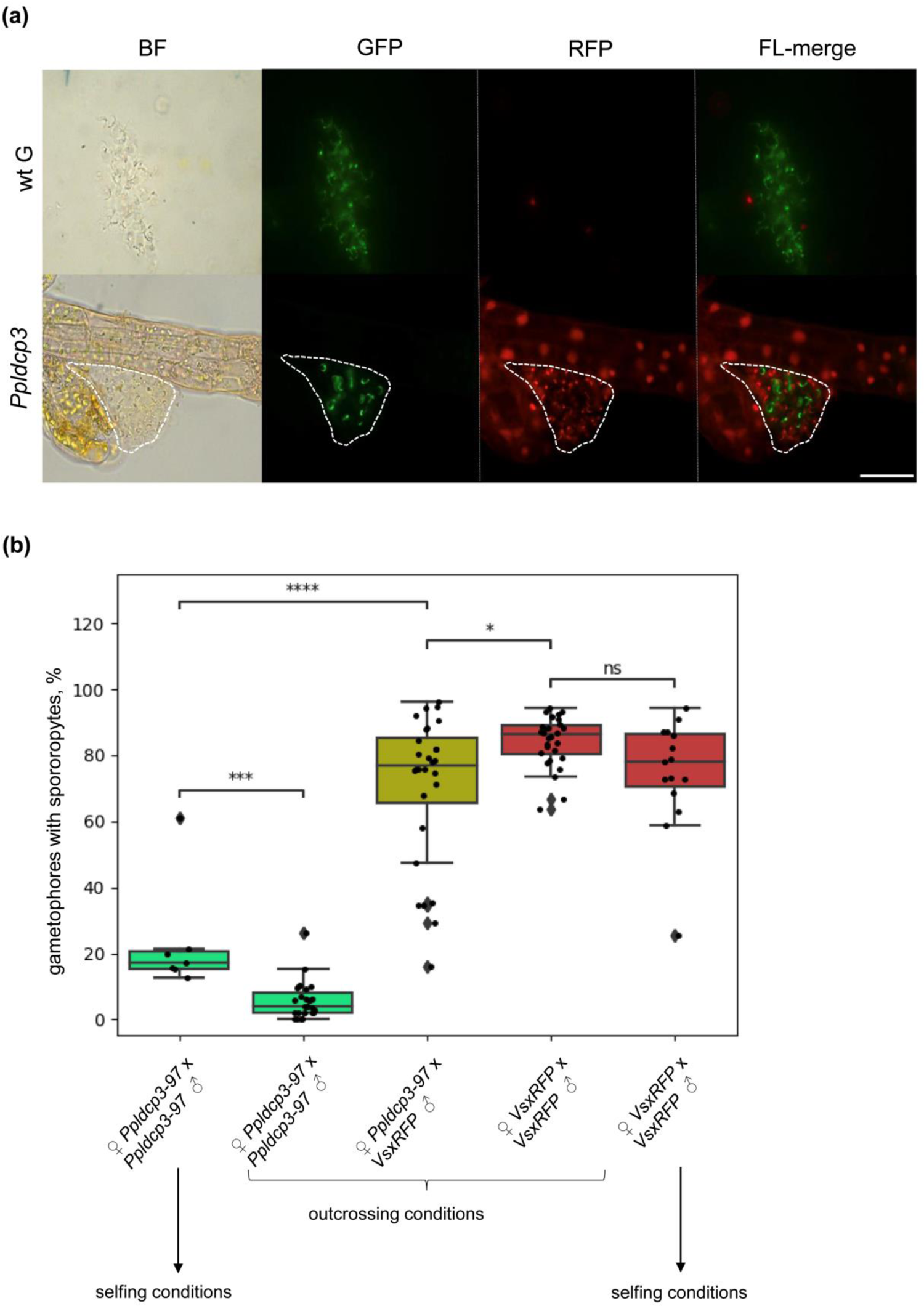
The Ppldcp3 phenotype in the gametophytic generation is paternal (a) Live-dead staining of antherozoids revealed two distinct populations of sperm cells. In the *Ppldcp3-97* mutant, a significant proportion of antherozoids was stained with propidium iodide (red channel), an indicator of compromised cell membranes of dead cells. In contrast, nearly all wild-type antherozoids were viable, specifically stained with SYTO 9 (green channel). Sperm release from antheridia in mutants was a rare event, with sperm being retained in the sperm matrix (shown in the image with a dashed line) for a long time after release. (b) Number of sporophytes produced in cross-fertilization of *Ppldcp3-97* mutants with highly fertile VsxRFP line. The majority of Ppldcp3 mutant shoots developed fluorescent sporophytes, supporting the hypothesis of a male reproductive defect being responsible for the reduction of sporophytes’ production on Ppldcp3 selfing plants. Statistical inference was performed using t-test independent samples with Bonferroni correction.

To further support and quantify antherozoid defects, we mixed protonemata of *Ppldcp3* and a highly fertile Villersexel line, constitutively expressing RFP protein under a maize ubiquitin promoter (*VsxRFP*) (Perroud *et al*., 2011), and grew them on soil. Supplementation of water on top of 15 dpi gametophores facilitates cross-fertilization between genotypically different shoots. The results of the experiment (Fig. 3b) revealed a high capacity for fertilization of *Ppldcp3* by *VsxRFP* sperm, distinguished as fluorescent sporophytes of nonfluorescent gametophores. The fertilization rates of *Ppldcp3-97* by *VsxRFP* were similar to those of selfing *VsxRFP* plants. Moreover, the number of self-fertilized *Ppldcp3* plants in outcrossing conditions (i.e. in competition with *VsxRFP* sperm) was significantly lower than in selfing conditions. These observations additionally prove that *Ppldcp3* sperm is mostly non-viable.

Multiple (up to 7) sporophytes often developed on a single *Ppldcp3* shoot in cross-fertilization conditions (Fig. S3a, b). All these sporophytes were fluorescent, thus being derived from multiple fertilization events on a *Ppldcp3* gametoecium and mediated by *VsxRFP* sperm. No more than 2 sporophytes were detected on self-fertilized *VsxRFP* plants.

All cross-fertilization-derived sporophytes, originating from single or multiple fertilization events on a shoot, developed normally showing neither abortions nor abnormal sporophytes. However, they were distinguishable from wild-type ones by larger size than either *VsxRFP* or *Ppldcp3-97* mutant sporophytes. No defects in spore release from these sporophytes were detected, and the spores developed normally and germinated similarly to the *VsxRFP* rates. Protonema derived from (♀ *Ppldcp3 x VsxRFP* ♂) germinated spores developed much faster than protonemata of *VsxRFP* and *wt Gransden* (Fig. S3c); an observation that might be explained by heterosis.

### PpLDCP3 localizes to male and female gametangia and is present in P-body-like structures

To assess the localization of PpLDCP3 in moss gametangia, we created lines expressing PpLDCP3 translationally fused with mCherry and mEGFP. Both PpLDCP3-mCherry and PpLDCP3-mEGFP C-terminal fusion lines were generated by knock-in in the Ppldcp3 locus, preserving the native promoter (Figs S4a, b). From the pool of candidates, three lines of each genotype were selected after verification by PCR and capillary sequencing, conferring the correctness of the homologous recombination (Fig S4c). Subsequently, PpLDCP3-mCherry-69 (PpLDCP3-mCh) and PpLDCP3-mEGFP-5 (PpLDCP3-mEGFP) were selected for imaging based on fluorescence intensity criteria in gametangia (Fig. S4d).

The fluorescent lines (Fig. S5a, see also Figs 4b, c), did not reveal fluorescent signal in intact vegetative tissue (leaflets observed), although a signal might be attributed to a particular cell type/ cell population in gametophores or associated with the regeneration of damaged tissue. Both epifluorescence and confocal microscopy for PpLDCP3-mCh (Fig. 4a) and PpLDCP3-mEGFP (Fig. S5a) revealed a relatively weak signal in antheridia that was specifically assigned to jacket cells, tip, and stalk antheridial cells. No fluorescence was detected by confocal microscopy in sperm cells, in agreement with transcriptomics data (Fig. S5b).

**Fig. 4.**
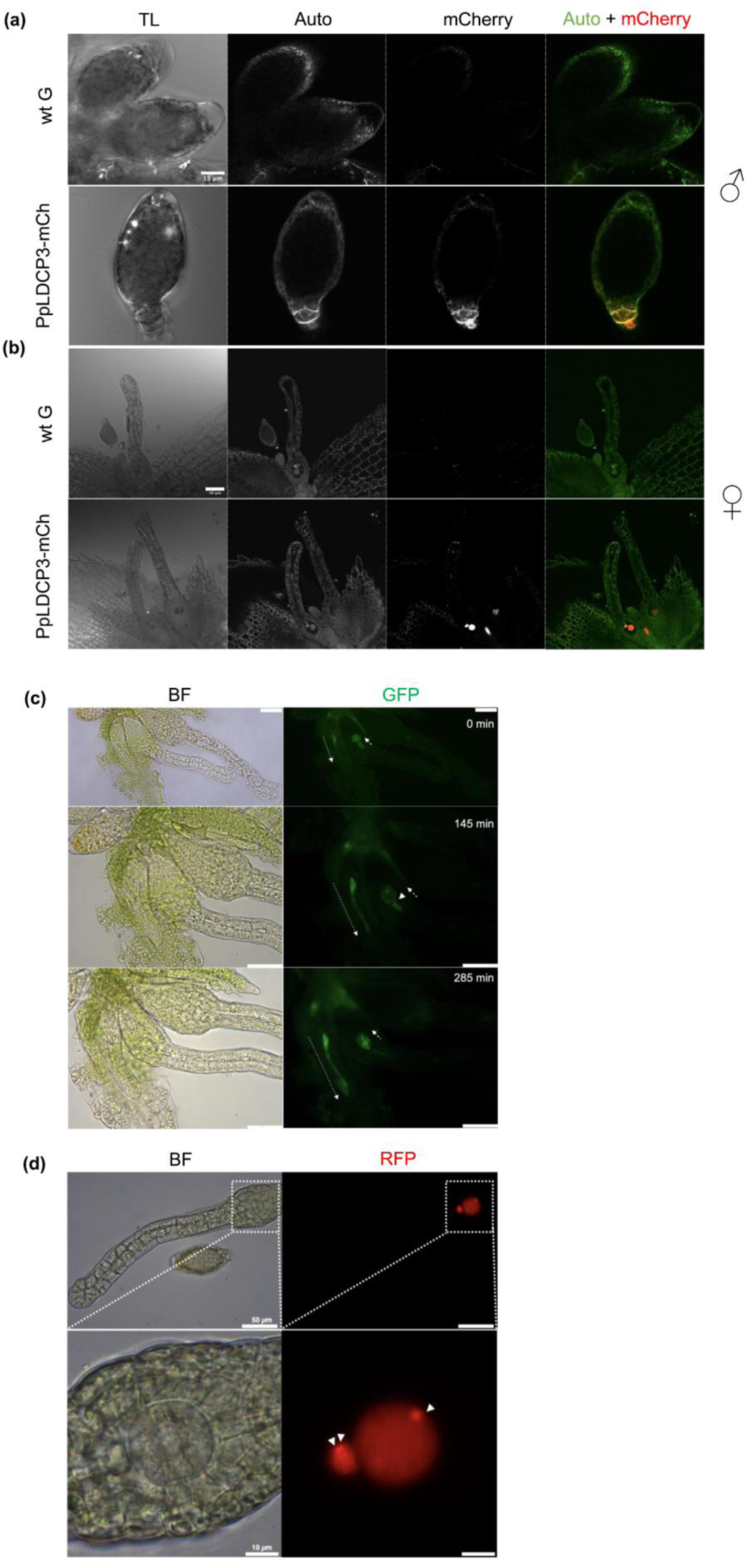
PpLDCP3-mCherry and PpLDCP3-mEGFP protein fusions localize to diverse cell types in developing gametangia (a) A weak autofluorescence signal was detected in the antheridia of PpLDCP3-mCh lines. The signal was localized to jacket cells, with the highest intensity at the antheridium stalk and tip cells. TL – transmitted light. (b) Confocal microscopy revealed the presence of PpLDCP3-mCh in archegonia. In young archegonia the signal is localised to canal and egg precursor cells (bottom panel, central archegonium). At the late stages of archegonium development, before the opening of the archegonium tip, the signal was localized to the egg cell and ventral canal cell (bottom panel, leftmost archegonium). After the degradation of the ventral canal cell, the signal from the egg cell disappeared before or right after the opening of the tip (bottom panel, rightmost archegonium). No signal was detected in intact leaflets or other parts of the gametophore body. (c) Time lapse of PpLDCP3-mEGFP localization in developing archegonia. The increase of the PpLDCP3-mEGFP signal over time during archegonium growth was accompanied by the elongation of canal cells (leftmost archegonium in all panels, the direction of fluorescent signal spread is shown with a dotted line arrow). The direction of fluorescent signal decline in the central archegonium, terminating its development, is shown with a dashed line arrow. At 145 minutes after the start of observation (middle panels) the ventral canal cell of the leftmost archegonium translocated at least part of its LDCP3-mEGFP containing cytoplasm to the egg cell, apparently, through the cytoplasmatic bridge (arrowhead). On the bottom panels, the signal in the central archegonium is detectable only in the egg cell. In all the panels, the rightmost archegonium is mature with an opened tip. No PpLDCP3-mEGFP signal was detectable in the rightmost archegonium during the time lapse observation. The scale bar is 50 µm. (d) Distinctive foci of PpLDCP3-mCh were observed in both egg and ventral canal cells of premature archegonia. The peripheral cytoplasmic foci (marked with white arrowheads) are visible before the degradation of the ventral canal cell or its partial fusion with the egg cell and are not observed in other canal cells. (see also this figure, c). The scale bar is 50 µm for the upper panel and 10 µm for the bottom panel. BF – bright field.

Strikingly, a strong fluorescent signal was revealed in developing archegonia (Fig. 4b). The signal localization had three spatiotemporal characteristics evolving during a few hours of observation (Fig. 4c, top to bottom): 1) localization is restricted to egg cell precursor and canal cells, with anterograde amplification throughout the archegonium neck, together with neck and canal cells expansion during organ development; 2) retrograde amplification of signal upon the maturation of the archegonium with localization to ventral canal cell and egg cell in premature archegonia (apex is closed); 3) transfer of PpLDCP3, apparently, through cytoplasmatic bridges from ventral canal cell to egg cell (apex is closed); 4) fading of signal in mature archegonia with opened apex (see Fig. 4b). Aging archegonia, with the canal turning brown, did not reveal any signal. PpLDCP3 in archegonial cells was found throughout the cytoplasm, however, the presence of the signal in the nuclei seemed to be more pronounced at the later stages of archegonium development. This agrees with the ubiquitous nature of PpLDCP3 localization predicted bioinformatically (Fig. S5d).

Additionally, a few intense foci were detected in maturing egg cells and ventral canal cells (Figs. 4d, S5c). Those might be similar to P-bodies, as described in mammalian oocytes (Flemr *et al*., 2010) or during meiotic exit in Arabidopsis (Cairo *et al*., 2022). We cannot exclude that these granules might result from protein misfolding and/or mislocalization, however, the general pattern of Ppldcp3 expression with assignment to egg cell can be confirmed by independent sequencing data (Sanchez-Vera *et al*., 2022) (Fig. S5e, f). Moreover, the presence of heat-response regions and extremely high AT-content in the promoter region of the Ppldcp3 gene (Fig. S5f) might suggest its role in the stress response and formation of stress granules in addition to P-bodies.

The effect of translational fusion on protein structure was hard to predict based on the modelling of the PpLDCP3 structure, as the predictions generated by AlphaFold show a very low level of confidence (Fig. S5g) that might be caused by intrinsically disordered regions (Fig. S5h). The latter is a common feature of LDCPs (Kubíková *et al*., 2021; Cipriani *et al*., 2021) and, at least to some extent, of many phase-separated proteins in general (Martin & Holehouse, 2020).

The accumulation of protein in the archegonium with no apparent functional female organ defect in the *Ppldcp3* mutant might be explained by the compensatory function of other proteins with similar domain architecture. While no direct homologs of PpLDCP3 were found in the *P. patens* proteome, several proteins with a similar domain set are present and expressed in egg cells, albeit at lower levels than PpLDCP3 (see Fig. S5e).

### Insights into molecular functions of PpLDCP3

To estimate the possible molecular and cellular function of PpLDCP3, we conducted bioinformatic predictions of its interaction partners. The STRING database assigned PpLDCP3 functionality to RNA metabolism, with multiple protein interaction partners involved in mRNA processing (Fig. 5a). These factors include polyadenylation and mRNA processing revealing parallelisms with animals’ LDCPs involvement in similar processes.

**Fig. 5.**
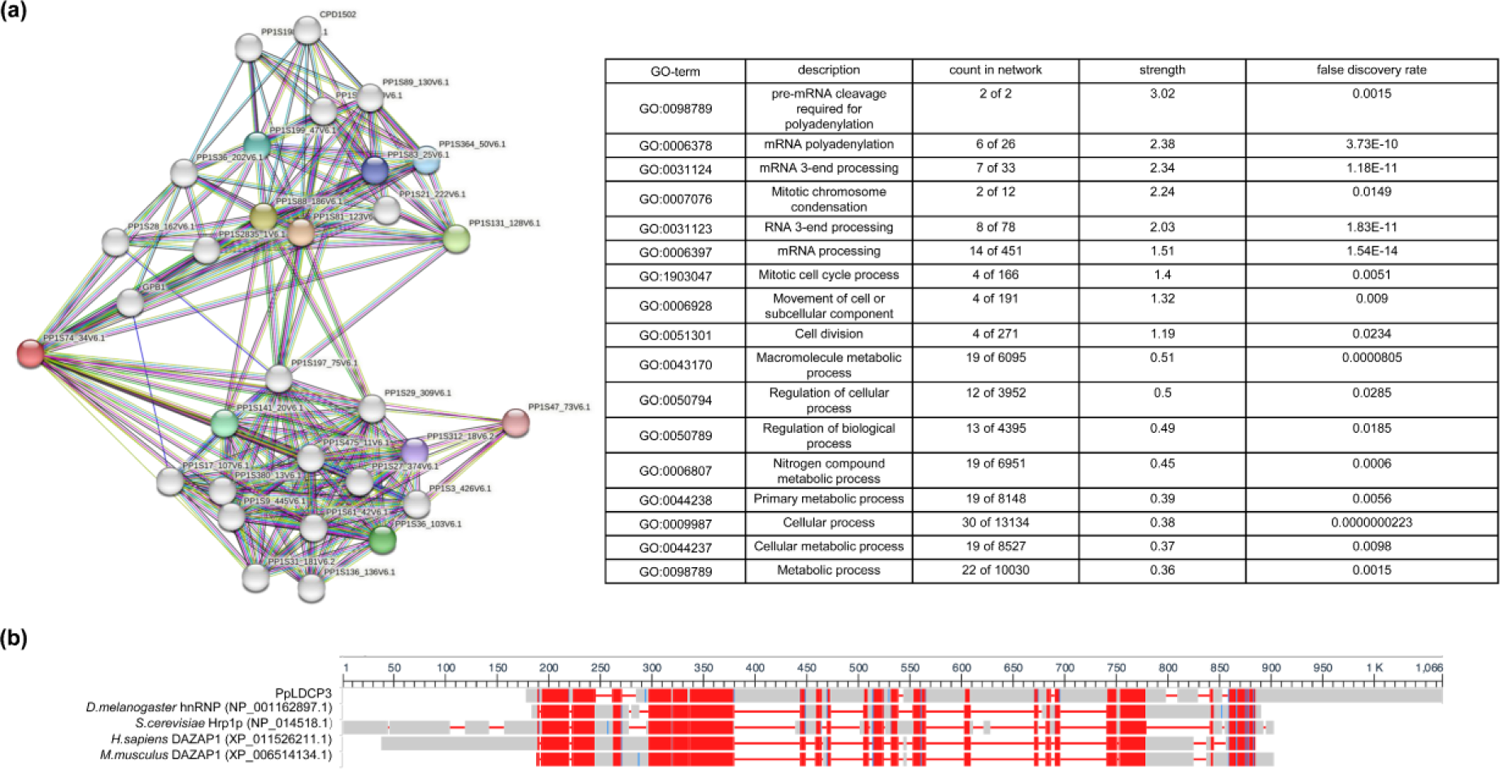
Insights into potential molecular functions of PpLDCP3 (a) Predicted interaction network for PpLDCP3 (Pp1s74_34V6.1; older community-based annotation) revealed by the STRING database. Interactions for 30 nodes are shown. The corresponding table shows Gene Ontology annotation based on the predicted nodes. (b) Remote homologs of PpLDCP3 outside of plant groups, revealed by SmartBlast and realigned by COBALT. Red blocks represent regions with a high level of similarity to PpLDCP3.

Another predicted functional group is associated with the mitotic cell cycle and cell division, providing further evidence for the possible involvement of PpLDCP3 in the synchronization of cell divisions. Cell cycle control by animal MARF1 is achieved through the control of phosphatases and cyclins, which in turn activate cyclin-dependent kinases. Serine residues, which are known to be the common phosphorylation sites, are predominant in PpLDCP3 (Fig. S5i). This amino acid content bias might reflect the existence of a feedback loop mechanism in the regulation of the cell cycle by PpLDCP3.

The molecular effectors related to the formation of functional sperm cells are better studied in animals. Alignment of PpLDCP3 with non-plant homologs (Fig. 5b) allowed us to establish distant similarity with RNA-binding proteins, among them heterogeneous nuclear ribonucleoproteins (hnRNP). The mammalian hnRNP DAZAP1 is the most similar protein to PpLDCP3 in mice and humans.

The appearance of the PpLDCP3 domain architecture in plants contributing both to the development of multicellular antheridium and sporophyte, required integration into the preexisting genetic networks controlled by master regulators of development. *A. thaliana* AT2G05160 shows similarities to PpLDCP3 domain architecture (see Fig. S5e) and was predicted to interact with WOX13 and bHLH (Fig. S5j), known to be crucial for plant development (Sakakibara *et al*., 2014; Lopez-Obando *et al*., 2022). Multiple WOX13 and bHLH binding motifs were found present in the promoter region of the Ppldcp3 gene (Table S1). Another predicted physical interactor, integrase, might be a link to the common function of animal LDCPs in transposon silencing during dispersal unit formation.

## Discussion

The ability to spread genetic material is essential for a species’ survival. Plant dispersal units play a crucial role in enabling the colonization of new habitats and facilitation of the gene flow between distant populations (Nathan, 2006; García *et al*., 2017; Rodger *et al*., 2018). Bryophyte ancestors have been the first to develop outer cellular layers of antheridia and sporophyte enclosing sperm and spores (Becker & Marin, 2009; Hafidh & Honys, 2021). These layers of cells might serve as nursing tissues and facilitate the timely maturation of propagules contributing to the ability to sense environmental cues appropriate for dehiscence (Paolillo, 1977; Hausmann & Paolillo, 1978; Brown & Lemmon, 2011; Lopez-Obando *et al*., 2022). It is tentative to speculate that the maturation and release of both types of dispersal units in moss might be regulated by common core gene players, thus coordinating the population’s success in an intertwined manner.

Here we have shown that the Ppldcp3 gene, encoding a LOTUS domain-containing protein 3, is expressed at the late stages of gametangia and sporophyte development and affects specifically dispersal units’ formation and dehiscence in *P. patens*.

### Ppldcp3 affects sporophyte development and spore dispersal

Sporophyte morphology is an important physiological and species trait in bryophytes (Rose *et al*., 2016). *Ppldcp3* mutants’ premature sporophytes often lost radial symmetry and appeared to have multiple developmental abnormalities. Dark-brown, mature *Ppldcp3* sporophytes contained highly vacuolized columella and amphithecium cells, the features of green sporophytes (stages 12-13, according to the development scheme provided in (Lopez-Obando *et al*., 2022)). However, 60-70 dpi *Ppldcp3* sporophytes contain mature-looking sporopollenin-covered spores, which are significantly smaller than wild-type ones.

Histochemical observations of *Ppldcp3-97* indicate defective sporogenesis marked by abnormal release of the spores from the tetrads, which might also explain the population of abortive cells in sporophyte preparations. In our experimental setting, *Ppldcp3* spores did not germinate. However, we could not fully exclude the viability of spores coming from a small proportion of sporophytes, considering the heterogeneity of the phenotypes produced by Pppldcp3 plants at all levels.

### Ppldcp3 is important for antheridia development and sperm maturation

By observing PpLDCP3 translationally fused with eGFP/mCherry we confirmed the expression of PpLDCP3 in the terminal stages of antheridia development. The protein was found localized to antheridial jacket cells, stalk, tip cells, and spermatocytes. Both imaging and transcriptomics data indicated the absence of PpLDCP3 expression in mature sperm. Similarly to the spores, *Ppldcp3* sperm were rarely found to be fully released. Sperm after release were retained within the sperm lipid matrix, with only part of the sperm appearing to be viable in live-dead staining. Fertilization incompetence of *Ppldcp3* sperm was confirmed by cross-fertilization with a highly fertile *VsxRFP* line.

The sum of the observations points to a complex male phenotype affecting antheridial development and sperm formation. The male *Ppldcp3* phenotype might be explained by the distant similarity of PpLDCP3 to DAZAP1 detected by SmartBLAST search, although no LOTUS domains are detectable in DAZAP1. DAZAP1 acts as a splicing activator (Choudhury *et al*., 2014), and its deletion leads to sterility in mice, with spermatogenesis disrupted in the pachytene stage of meiosis (Hsu *et al*., 2008). This might denote to a highly conserved role of the LDCP3 protein sequence pattern in evolution of motile gamete formation.

### PpLDCP3 in maturing archegonia may be involved in hormonal-mediated signalling

Lines expressing PpLDCP3 fused with fluorescent proteins revealed a strong signal in the female germline, namely in archegonia canal cells and premature egg cells. However, the role of PpLDCP3 in archegonia seems different from the roles observed in antheridia or sporophytes. This is supported by the fact that most of the *Ppldcp3* shoots appeared to be fertilized by *VsxRFP* sperm. Moreover, *Ppldcp3* archegonia acquire much higher fertilization competence, resulting in multiple independent fertilizations occurring on the same *Ppldcp3* shoot concurrently. This observation might resemble the phenomenon of superfecundation in animals (Jonczyk, 2015) and is similar to multiple fertilizations taking place in more evolutionary advanced embryophyte groups. The increase in the archegonia number and/or simultaneous readiness of multiple eggs to be fertilized might be the physiological explanation for the observed phenotype. It is possible that the lack of Ppldcp3 results in a compensatory effect from proteins with similar domain architecture, ultimately leading to hyperactivation of the female germline.

Several genetic factors, among which are auxin transporters PppinB and PppinC (Bennett *et al*., 2014; Lüth *et al*., 2023) are known to restrict the number of fertilized archegonia, and, thus, may limit the sporophyte production to a single one per shoot. This connection of PpLDCP3 with auxin signaling seems a viable explanation as a mutant of the regulator of auxin biosynthesis, PpSHI2, demonstrates problems with sperm release from antheridia, another link to PpLDCP3 function in male germline. These pieces of evidence may indicate broader roles of a putative auxin-transporters - LDCP crosstalk, affecting different stages of gametangia development.

### PpLDCP3 translational fusion lines reveal aspects of archegonia development

The observation of PpLDCP3 protein localization using reporter lines also provided valuable insights into germline development in *P. patens*. During archegonia maturation, the degradation of the canal cells is followed by the disappearance of the fluorescence signal in the archegonium neck. In the subsequent stages, at least part of PpLDCP3 containing cytoplasm seems to translocate from the ventral canal cell to the egg cell. This translocation seems to occur through cytoplasmatic bridge-like structures and is followed by the concentration of the signal in the egg cell, with its subsequent fading correlated with the archegonium opening.

The functions of the PpLDCP3 redistribution in the archegonia cells are not clear, but it appears reminiscent of the translocation of molecules from nursing cells to gametes in the developing animal germline (Mische *et al*., 2007). In plants, symplastic communication between the egg and the central cell was shown for *Torenia fournieri* (Han *et al*., 2000). Similarly, the translocation of polypeptides and small RNAs has been investigated in the *Arabidopsis* female germline (Schröder *et al*., 2022). To our knowledge, no evidence of germline cell communication has been shown in *P. patens*.

### PpLDCP3 might share common functions with animal LDCPs related to RNA metabolism

The observed LDCP3-mCherry and LDCP3-mEGFP fluorescent foci in archegonial cells point to a role of PpLDCP3 as a player in biomolecular condensates. LDCPs, such as TDRD5/7, Oskar, MIP1/2, and LOTR-1 are known to be localized in germ granules of either developing gametes or supporting nurse cells of male and female animal gonads (Yabuta *et al*., 2011; Tanaka *et al*., 2011; Ding *et al*., 2018; Kubíková *et al*., 2021; Pamula & Lehmann, 2024).

However, in plant gametogenesis and sporogenesis, no specific germ granules have been described and limited evidence exists for the involvement of biomolecular condensates in plant reproduction (Oliver & Martinez, 2022; Cairo *et al*., 2022). As stress granules and processing bodies are the only two types of biomolecular condensates described in the plant cytoplasm (Emenecker *et al*., 2020; Kearly *et al*., 2022), we refer to the observed condensates as P-bodies. However, both P-bodies and stress granules share certain similarities in functionality and composition (Banani *et al*., 2017) and the detection of heat-response elements in the 5’-region of the Ppldcp3 gene and low-melting temperature sequence in the promoter region of the gene might be evidence for the protein activation additionally in stress-related conditions.

In P-bodies, LDCPs may participate in RNA decay, as it was shown for MARF1 in human cell lines and DNE1 in *A. thaliana* vegetative tissues (Nagarajan *et al*., 2023). Ribonuclease activity was also described for a group of mitochondrial LDCPs, MNU1 and MNU2 (Stoll & Binder, 2016) from *A. thaliana*. However, PpLDCP3 lacks a detectable ribonuclease domain. On the other hand, an *A. thaliana* protein encoded by the AT2G05160 locus possesses the same domain set as PpLDCP3, and demonstrates ribonuclease activity (Addepalli & Hunt, 2008). An intrinsic ribonuclease activity of PpLDCP3 might therefore be worth investigating.

Mutants of animal LDCPs have been characterized by deregulation of the germline cell cycle interfering with cyclin turnover, which in turn affects cyclin-dependent (de)phosphorylation (Su *et al*., 2012; Kawaguchi *et al*., 2020). High abundance of potentially phosphorylated serine in the PpLDCP3 protein sequence together with histochemical evidence of abnormal cell divisions in the *Ppldcp3* sporophyte may serve as indicators of regulatory loops, controlling cell cycle progression mediated by PpLDCPs. The potential role of PpLDCP3 as a regulator of cell division needs further investigation.

One predicted interactor of AT2G05160-derived LDCP is an integrase, belonging to an enzyme family that manages retrotransposition of mobile genome elements. In animal LDCP mutants, germline transposon activation typically occurs (Patil & Kai, 2010; Tanaka *et al*., 2011; Su *et al*., 2012; Patil *et al*., 2014). In plant germlines activation of developmentally relaxed transposable elements was shown (Slotkin *et al*., 2009; Martínez & Slotkin, 2012; Long *et al*., 2021), which, when uncontrolled, may lead to gene disruption due to TE insertion. It is possible that the absence of PpLDCP3’s safeguarding function in *Ppldcp3* mutants may lead to transposon integration into the germline genome resulting in sperm inviability and subsequently abnormal sporophyte shapes.

### PpLDCP3-like domain architecture might be important for the formation of plant adaptations to terrestrial environments

Despite the absence of detectable protein similarity, the PpLDCP3 domain set is the same as in *A. thaliana* proteins encoded by AT3G52980 and AT2G05160 loci. We suggest that these proteins, along with their homologs in other plant groups, might be considered plant germline LDCPs. Similar to PpLDCP3, the proteins derived from AT3G52980 and AT2G05160 loci share germline expression patterns and contain Zinc-finger domains of the C3H1 and/or C2H2 type, which are likely plant-specific architecture novelties for LDCPs. Zinc-finger proteins are known for their versatile binding to DNA, RNA, and proteins, influencing plant growth, stress responses, vegetative tissue formation, fruit ripening, and seed germination (Bogamuwa & Jang, 2013, 2014; Liu *et al*., 2022). The PpLDCP3 structural novelty might be responsible for versatile molecular interactions in the terminal developmental stages of diverse dispersal units.

Efficient spore dehiscence is crucial for dispersal (Johansson *et al*., 2014), but mature spore capsules rarely open in *Ppldcp3*. Similarly, sperm release was perturbed in *Ppldcp3* lines. These defects might be due to perturbation of the networks related to bHLH, which is predicted to interact with AT2G05160-derived protein in *A. thaliana*. In angiosperms, bHLHs are crucial for pollen and seed development, their dehiscence and viability (Qi *et al*., 2015; Walbot & Egger, 2016; Wei *et al*., 2018). In *P. patens*, bHLH proteins are responsible for spore production, providing nursing functions for sporocytes (Lopez-Obando *et al*., 2022).

Another predicted interactor of the protein encoded by AT2G05160, AtWOX13, is essential for seed dehiscence and the opening of the silique (Ferrándiz, 2002; Roeder *et al*., 2003; Romera-Branchat *et al*., 2013; Herrera-Ubaldo & de Folter, 2022). In PpLDCP3 mutants, fewer sporophytes and their abnormal shapes were observed, similar to mutants of PpWOX13L (Sakakibara *et al*., 2014).

Summing up these observations, we suggest that PpLDCP3 may function downstream of WUSCHEL and bHLH genes, controlling the final stages of cell speciation and tissue connectivity, and genome safeguarding, crucial for dispersal unit formation. Phase separation processes might be prerequisites for dispersal unit formation through RNA degradation, with PpLDCP3 serving as an integrative player in the process. Identifying pathways involving PpLDCP3-like proteins may help in understanding the evolution of land plant dispersal units and identify mechanisms to manipulate agriculturally important traits, such as fruit ripening and shattering (Li *et al*., 2006; Funatsuki *et al*., 2014).

## Acknowledgements

The authors are grateful to the IGC Plant Facility and especially to Vera Nunes. Sporophyte histological sample processing and staining was performed at the Histopathology Unit of the IGC. Pedro Carvalho (ITQB NOVA - Instituto de Tecnologia Química e Biológica António Xavier) instructed AK on the epifluorescent microscopy and helped with video recording. Confocal microscopy images were produced using the microscopes from the Bacterial Imaging Cluster (ITQB NOVA), members of which also provided training and guidance for microscope usage. Their work is partially supported by PPBI - Portuguese Platform of BioImaging (PPBI-POCI-01-0145-FEDER-022122) co-funded by national funds from OE - “Orçamento de Estado” and by European funds from FEDER - “Fundo Europeu de Desenvolvimento Regional”). We are grateful to M. Bezanilla’s lab for providing plasmids for construction translational fusion lines. This work was supported by FCT - Fundação para a Ciência e a Tecnologia, I.P., through GREEN-IT Bioresources for Sustainability” Unit (UIDB/04551/2020, DOI: 10.54499/UIDB/04551/2020), and through LS4FUTURE Associated Laboratory (LA/P/0087/2020; DOI: 10.54499/LA/P/0087/2020). JDB received salary support from FCT through grant CEECIND/03345/2018. AK received FCT scholarship PD/BD/128434/2017 and funding from Instituto Gulbenkian de Ciência.

## Competing interests

The authors declare no competing interests.

## Author contributions

AK and JDB conceived the idea. AK performed the experiments, prepared figures, and wrote the manuscript. JDB revised figures and manuscript.

## Data availability

Scripts for data analysis are available on GitHub (https://github.com/avkermanov) and can be provided upon request.

## Supporting Information

### Supplementary methods

Supplementary material and methods not included in the main text

**Figure S1:** Validation of *Ppldcp3* mutants and phenotypic effects related to sporophyte development

**Figure S2:** Quantification of spore sizes in the wild-type and *Ppldcp3* mutant

**Figure S3:** Hybrid sporophytes develop on *Ppldc3* gametophores in cross-fertilization experiments with the *VsxRFP* line

**Figure S4:** Generation and verification of PpLDCP3 translational fusion lines

**Figure S5:** Localization of PpLDCP3 in gametangia and its predicted genetic and biochemical properties, responsible for germline and dispersal unit formation

**Video S1:** *Live* Ppldcp3 sporophyte dissection

**Table S1:** Predicted binding sites for WOX13 and bHLH transcription factors in the promotor region of the Ppldcp3 gene

**Table S2:** Primers used in this study

## Notes

### Competing Interest Statement

The authors have declared no competing interest.

