## Supplemental Methods for "The LOTUS-domain containing protein PpLDCP3 controls germline and dispersal unit formation in the moss *Physcomitrium patens*"

### Supplementary methods

#### Plant growth on soil

Spores or ground protonema were placed on autoclaved and rehydrated peat pellets (Jiffy-7; Jiffy Products) (referred to in the text as “jiffies”). For cross-fertilization experiments, equivalent amounts of ground protonemata were mixed before placing on the soil. Plants on pellets were grown in transparent plastic boxes (4-5 jiffies per box), regularly filled with water up to half of the box, under controlled conditions of a “long day”: 16 h light (light intensity 90 μmol/($m^2$s)), 25 °C, and 50% humidity. To induce reproductive organ formation, plants were first grown under long-day conditions for 4-6 weeks and were then transferred to a “short day” chamber with the following conditions: 8 hours of light (light intensity of 80 μmol/($m^2$s), 17°C, and 50% humidity. The induction was verified by stereoscopic observation of reproductive organs’ formation on 10-15 dpi shoots. In the cases in which water supplementation was a part of the experimental setting, 30 ml of sterile water was supplemented drop-wise using a 20 ml glass pipette on top of gametophores at 15,17, and 19 days past induction (dpi) to facilitate fertilization.

#### Plant DNA extraction, amplification of fragments for construct generation, and genotyping

*DNA extraction. P. patens* gDNA was extracted from 5–7 day-old protonema or developing gametophores before induction. Plant tissue was first frozen in liquid nitrogen and then ground with a disposable pellet pestle in TE buffer. Further extraction was performed with a GRISP plant gDNA extraction kit according to manufacturer instructions. The concentration and purity of extracted DNA were measured with Nanodrop™ 2000 and approximately 1ng of gDNA was used in PCR reactions.

*Plant DNA amplification for molecular cloning and genotyping:* Genotyping PCR reactions were performed in 10-20 ul volume using a KAPA 3G Plant PCR kit or a DreamTaq DNA polymerase kit with designated primers according to standard conditions. The amplified PCR product was verified by running 1-2 % TAE agarose gels. Amplifications for molecular cloning were performed using Phusion DNA polymerase or AccuTaq DNA polymerase directly from genomic DNA or after preamplification (5-7 cycles) with a KAPA 3G Plant PCR kit.

**Bacterial plasmid DNA extraction and construct verification**

Mach1 or DH5α chemically competent cells, prepared in-house using Mix & Go! *E. coli* Transformation Kit and Buffer Set (Zymo), were used for transformations. Selection of positive colonies was performed on LB agar medium, supplemented with ampicillin to 100 µg/ml. Selected colonies were grown in 3 – 10 ml of liquid LB medium with the addition of ampicillin to 100 µg/ml. Plasmid DNA was extracted using the GRS Plasmid Purification Kit – Mini kit, following the manufacturer's instructions. Verification of plasmid inserts was done by PCR using DreamTaq DNA polymerase, by analytic digestion with a panel of restriction enzymes, and by capillary sequencing.

**Generation of constructs for plant transformation and for obtaining mutant plant lines**

*Ppldcp3 full knock-out mutant construct generation.* pBHRf vector was modified to carry the right and left flanks of Ppldcp3 gene, with a Hygromycin resistance cassette in between, as described elsewhere (Schaefer *et al.*, 2010). Briefly, left and right homologous arms were amplified in the reactions with AK63+AK70 and AK64+AK65 primers. Each arm was introduced into the pCR4-TOPO plasmid. Next, the left arm was recovered from the plasmid and cloned into pBHRf, linearized with AvrII/PmeI restriction enzymes. The generated vector was linearized with AscI/HpaI restriction enzymes, and the right flank fragment was subcloned. The final vector was linearized with AvrII and AscI and used for wild-type Gransden protoplast transformation. Plasmid construction details are shown in supplementary Figure S1A.

*PpLDCP3 translational fusion construct generation.* Translational fusion lines were created using Gateway technology with a system with four-fragment recombination vectors and pGEM-gate as destination vectors, all kindly provided by M. Bezanilla's lab. C-terminal translational fusions were introduced by homologous recombination in the Ppldcp3 locus, removing the stop codon and fusing Ppldcp3 open reading frame sequence with mEGFP or mCherry coding sequences, deprived of the start codon. G418 resistance cassette was introduced next to the fluorescent protein coding sequence. Briefly, two regions up (left flank) and downstream (right flank) of the Ppldcp3 stop codon were amplified to be used as homologous arms for recombination. For the left flank insert, primers AK146 and AK147 were used to amplify the fragment, inserted then in pCR4-TOPO. The right flank was amplified using primers AK149 and AK150 and inserted into another pCR4-TOPO vector. Both modified vectors were amplified with primer pairs AK148 and C29 - for the left flank and AK151 and

C30 for the right flank to generate attB sites compatible with BP recombination reaction with pDONR. After BP recombination, the vectors carrying right and left flanks were recombined (LR reaction) with 2 other vectors, carrying mCherry/mEGFP and G418 resistance cassettes, resulting in the destination vector carrying 2 homologous arms neighboring Ppldcp3 (subject to removal), stop codon and both fluorescent protein-coding gene and G418 resistance in between the arms. Vectors were linearized with AvrII and PacI and used in the wild-type Gransden protoplast transformation. Plasmid construction details are described in supplementary Figures S4A, B.

**Plant transformation, selection of transformants, and verification of stable lines**

Plants were transformed according to the protocol (Schaefer *et al.*, 1991), with modifications. Briefly, for each transformation wild-type protonema of *P. patens* Gransden ecotype was cultivated on agar plates in the regular growth (long day) conditions as described above for at least 2 weeks and was ground at least twice during this period. 4-5 day-old protonema was collected from plates and subjected to driselase enzymatic protoplastation in 0.5 M mannitol. Protoplasts were separated from debris with a sieve, followed by low-speed (800 rpm, bucket rotor for 15 ml tubes) centrifugation. An aliquot was collected to count the round-shaped protoplasts with a hemocytometer and protoplast concentration was adjusted with 3M media to around ($1.2x10^6$ protoplasts)/(ml). To 300 µl of protoplasts, an equivalent volume of PEG-containing solution was added (40% PEG; 0.1 M Ca(NO3) in 3M medium). 15-30 µg of plasmid was used for each transformation. pBAS plasmid, containing a constitutively expressed GFP was used as transformation control. The transformation was performed by heat shock in a water bath at 45 C for 15 minutes. The osmotic pressure in the solution was restored by the gradual addition of a 3M medium to the transformation mixture. Then regeneration conditions were established by low-speed centrifugation followed by the gradual replacement of 3M medium with 3 ml of sugar-rich REG medium (5% glucose, 3 % mannitol in KNOPS medium) and transfer to a 30 mm Petri dish. Protoplasts were left to regenerate the cell wall in darkness for 5 days and then transferred to standard *P. patens* chamber long day conditions for the next 5-7 days. Regenerated protoplasts were plated on KNOPS-T plates containing antibiotics. Protoplasts that were able to regenerate and develop protonema under antibiotic-selective pressure for 20 days, were then transferred to an antibiotic-free medium. After 2-3 weeks plants were transferred back to Hygromycin or G418 medium to ensure the non-transient nature of acquired resistance. Finally, the surviving resistant plants were genotyped, and the insertion locus was verified by Sanger sequencing. The plants supported by sequencing evidence were selected to grow the full cycle on soil, and subsequently 2-3 stable lines were picked for further characterization.

**Spore counting and size measurement**

*Spore counting.* Sporophytes at the mature stages, but not ruptured, were collected from both wild-type and *Ppldcp3* mutant plants. The least abnormal mutant sporophytes (wt-type) were selected to avoid biased estimates. Sporophytes were fixed in FAA solution (formaldehyde: acetic acid: alcohol (1:1:18) and 3 random sporophytes were placed in an eppendorf tube, washed with water, and spores released in 100 ul of water. The solution was centrifuged at 94 g in a table centrifuge for 1 min, 75 ul of supernatant was removed and the pelleted spores were resuspended in 25 ul of water. 11 ul were placed on a hemocytometer chamber in 2 replicates and 4 squares corresponding to 0.1 ul volume were counted 3 times and averaged, generating one measurement.

*Calculations of spores' sizes.* Images of wild-type and *Ppldcp3* spores were converted to 8-bit, and the threshold was adjusted equivalently. The background was subtracted with a rolling ball radius set at 50. The image was processed to binary with filling holes and "watershed" applied to separate closely related objects. Spores' sizes on modified images were analyzed with the "analyze particles" tool. The preselected size was set (150-1000 $\mu m^2$) and particles corresponding to spores were selected on the initial image and then the areas of selected regions of interest were calculated automatically. The spore sizes on the graph represent at least 3 independent biological replicates.

**Histochemical sectioning and staining**

Sporophytes for histological sectioning were collected from wild-type and *Ppldcp3* plants at the late developmental stages and fixed in FAA (formaldehyde: acetic acid: alcohol (1:1:18)) solution. Further sample processing was performed at the Histopathology Unit of the IGC. Briefly, samples were embedded in 3% agarose and subjected to gradual dehydration in increasing concentrations of ethanol, (30% to absolute ethanol) at $4^{o}$ C with agitation for 2 days. Then sporophytes were pre-impregnated in Technovit 7100-ethanol (1:1) and left overnight. This was followed by 1 day of methacrylate impregnation mixed with Hardener I and followed by impregnation, embedding, and polymerization with methacrylate in Hardener II. Sectioning was performed using Leica RM2135, producing a series of 3 μm thin slices transversally and longitudinally. Slides were stained for 5 minutes in 1% TBO with 2% Sodium Borate and washed with PBS+T.

**Bioinformatics approaches for revealing additional Ppldcp3 features**

GC content of the Ppldcp3 gene and flanking areas were calculated by VectorBuilder (https://en.vectorbuilder.com/tool/gc-content-calculator.html) with a sliding window of 200 bp.

HRE elements (consensus sequence nTTCnnGAAn) (Xiao & Lis, 1988), WOX and bHLH binding sites (Toledo-Ortiz *et al.*, 2003; Ikeuchi *et al.*, 2022) were identified using the SnapGene tool. Protein localization in the nucleus was estimated with NucPred (Brameier *et al.*, 2007). The PpLDCP3-mCherry protein structure model was predicted *de novo* using AlphaFold Protein Structure Prediction (Jumper *et al.*, 2021) integrated into ChimeraX-1.5 (Goddard *et al.*, 2018; Pettersen *et al.*, 2021). Intrinsically disordered protein regions were predicted using IUPred3 (Mészáros *et al.*, 2018; Erdős & Dosztányi, 2020; Erdős *et al.*, 2021) with settings: "IUPred3 long disorder, medium smoothing". All predictions were done using the longest PpLDCP3 isoform. Prediction of the PpLDCP3 interactome was performed using STRING Interaction network version 11.5 (Szklarczyk *et al.*, 2019) generated for the Ppldcp3 gene. Physical interaction of the *A. thaliana* protein encoded by AT2G05160 was generated by ThaleMine v5.1.0-20221003 (Krishnakumar *et al.*, 2016).
