## Supplemental Figure S1 for "The LOTUS-domain containing protein PpLDCP3 controls germline and dispersal unit formation in the moss *Physcomitrium patens*"

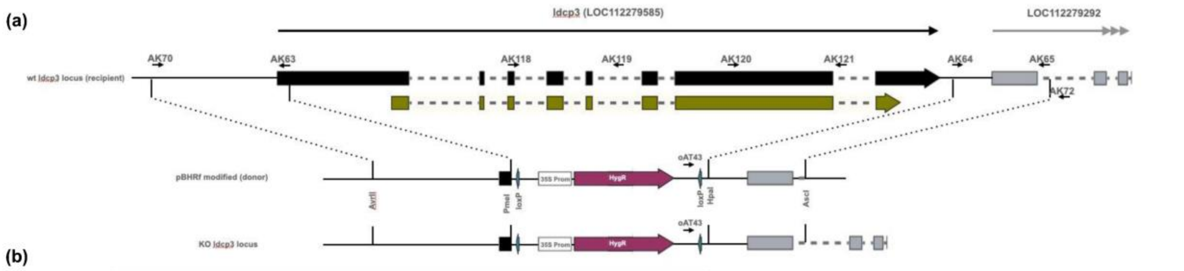


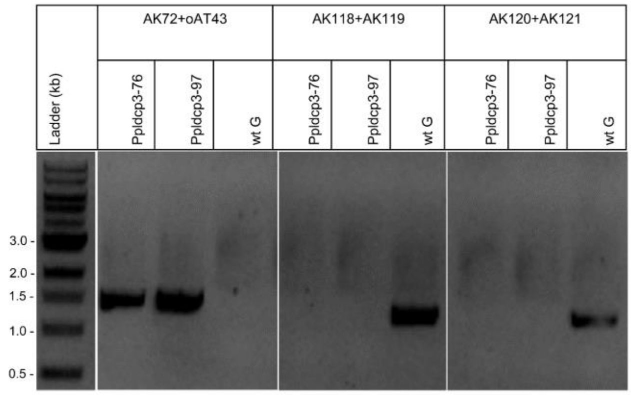


| Primer pair used in PCR | Amplicon length in wt G, bp | Amplicon length in *Ppldcp3*, bp |
|---|---|---|
| AK72+oAT43 | no amplification | 1374 |
| AK118+AK119 | 1238 | no amplification |
| AK120+AK121 | 1267 | no amplification |

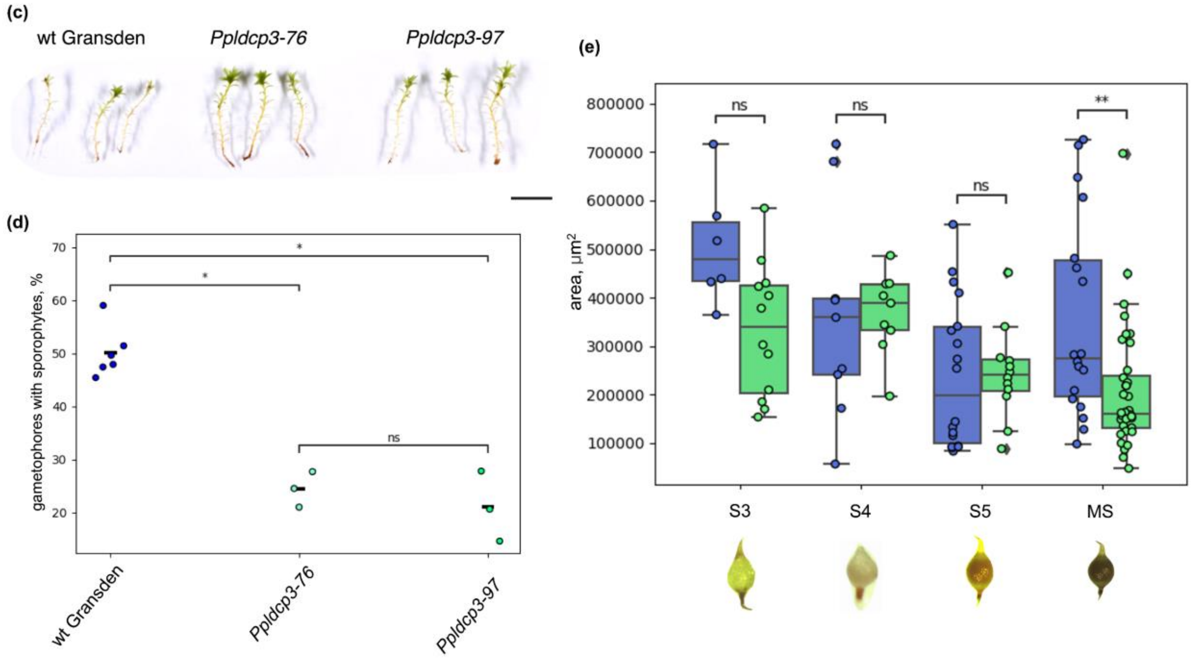


**Figure S1. Validation of *Ppldcp3* mutants and phenotypic effects related to sporophyte development**

(a) – Representation of Ppldcp3 locus with primers used in this study and *Ppldcp3* knock-out mutant generation scheme.

(b) – On the right panel the compilation of agarose gels is shown, confirming Ppldcp3 replacement with antibiotic resistance cassette for lines *Ppldcp3-76* and *Ppldcp3-97* (reaction AK72+oAT43). Additional PCR tests (AK118+AK119; AK120+AK121) were carried out to confirm the absence of the gene reintegration elsewhere in the genome. On the left panel, the expected length of the PCR products and the corresponding output of the amplification are presented in table format.

(c) – Developed gametophores of wild-type Gransden and *Ppldcp3* mutants were collected and imaged after growing on soil 50 days post-induction. No obvious developmental defects were observed during the vegetative growth of *Ppldcp3* mutants compared to wild-type plants. The scale bar is 5 mm.

(d) – Supplementation of water on top of gametophores on 15, 17, and 19 dpi results in a proportional increase of the number of mature sporophytes counted on 60-70 dpi. Statistical inference was performed using the Mann-Whitney-Wilcoxon test two-sided with Bonferroni correction.

(e) – Difference in sporophyte sizes from different developmental stages (meiotic-postmeiotic, green, premature, mature) of wild-type and *Ppldcp3-97* mutant lines, representing the significant reduction of sporophyte sizes of *Ppldcp3* mutant (green) compared to wild-type Gransden (blue). Statistical inference was performed with a t-test for independent samples with Bonferroni correction.
