## Supplemental Figure S2 for "The LOTUS-domain containing protein PpLDCP3 controls germline and dispersal unit formation in the moss *Physcomitrium patens*"

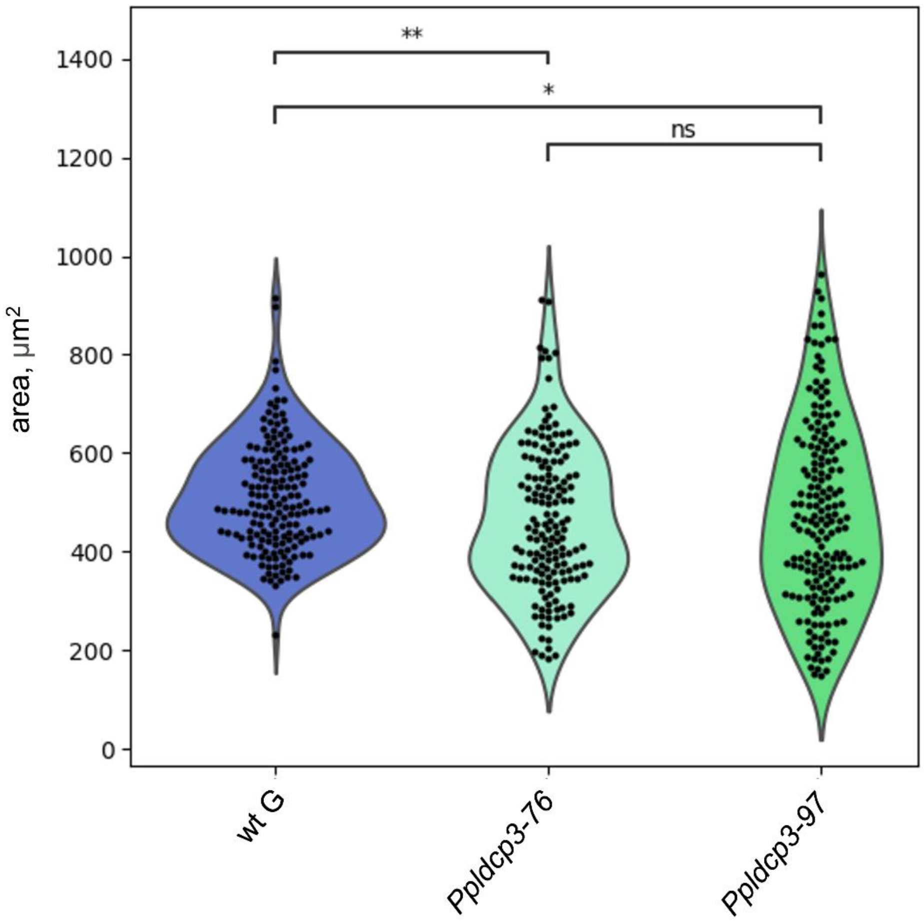


**Figure S2. Quantification of spore sizes in the wild-type and *Ppldcp3* mutant**

Only large spore-like particles were included in the counting in the mutant plants. The graph shows the overall smaller size and higher variation in size of the *Ppldcp3* spores compared to the wild-type. Statistical inference was performed using t-test independent samples with Bonferroni correction, where p is the P-value (adjusted).
