## Supplemental Figure S3 for "The LOTUS-domain containing protein PpLDCP3 controls germline and dispersal unit formation in the moss *Physcomitrium patens*"

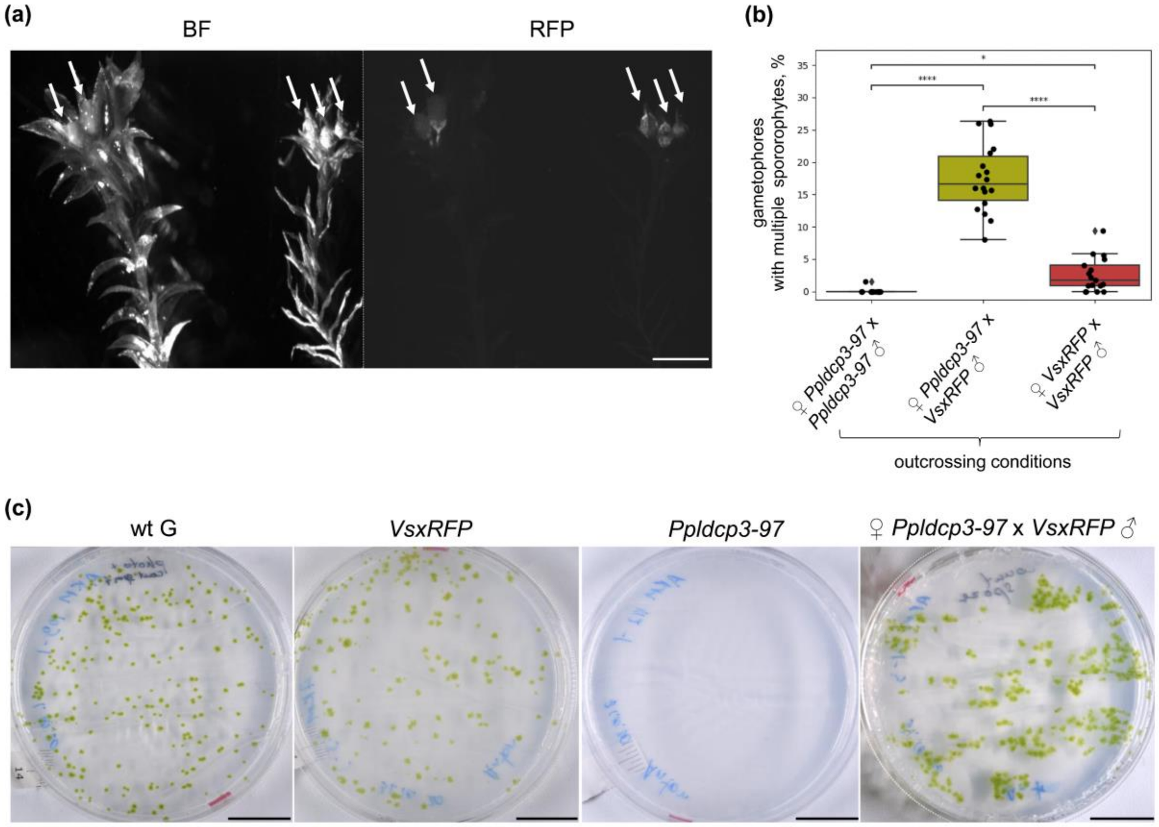


**Figure S3. Hybrid sporophytes develop on *Ppldc3* gametophores in cross-fertilization experiments with the *VsxRFP* line**

(a) – Multiple sporophytes on *Ppldcp3* shoots, resulting from cross-fertilization experiments with *VsxRFP* sperm (♀*Ppldcp3* x *VsxRFP*♂), were observed. Sporophytes arise separately from the same gametangia cluster, indicating the fertilization of multiple archegonia took place. All sporophytes (indicated with arrows) emit in the RFP fluorescent channel, pointing to fertilization by *VsxRFP* sperm. The scale bar is 5 mm.

(b) – The number of shoots with multiple sporophytes in outcrossing conditions was measured for different combinations at 60 dpi. All multiple sporophytes in ♀*Ppldcp3* x *VsxRFP*♂ cross were fluorescent. Statistical inference was performed using t-test independent samples with Bonferroni correction.

(c) - Hybrid (after cross-fertilization ♀*Ppldcp3* x *VsxRFP*♂) fluorescent sporophytes were collected to compare the germination of the spores in comparison with other genotypes. Spores of hybrid sporophytes were germinating successfully, visually comparable to wild-type rates, but developing protonema faster than either of the parental plants. The scale bar is 2 cm.
