## Supplemental Figure S4 for "The LOTUS-domain containing protein PpLDCP3 controls germline and dispersal unit formation in the moss *Physcomitrium patens*"

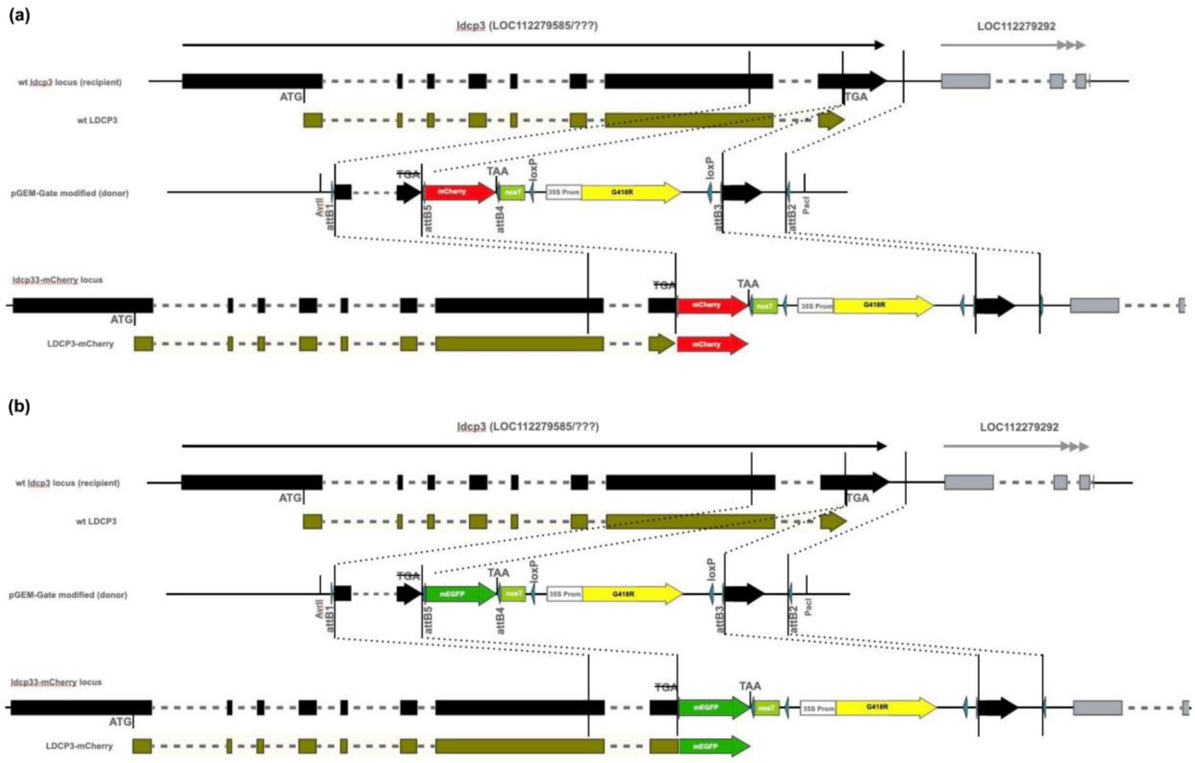


(c)

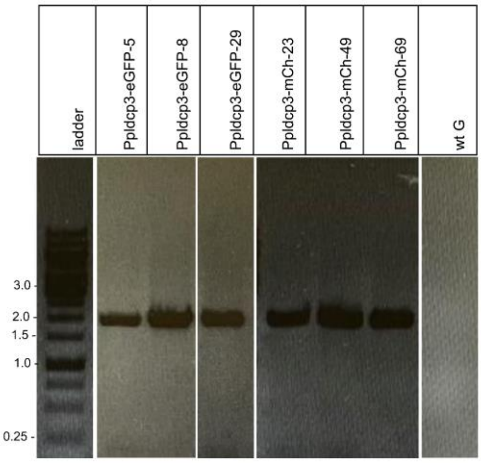


| Primer pair used in PCR | Amplicon length in wt G, bp | Amplicon length in Ppldcp3-mCherry | Amplicon length in Ppldcp3-eGFP |
|---|---|---|---|
| AL028+AK72 | no amplification | 1967 | 1967 |

(d)

| line | C-terminal protein, fused to LDCP3 | gametophores with sporangiophores, % | Fluorescent signal intensity |
|---|---|---|---|
| Ppldcp3-mCh-69 | mCherry | 12.4 ± 2.0 | +++ |
| Ppldcp3-mCh-23 | mCherry | 17.2 ± 0.9 | + |
| Ppldcp3-mCh-49 | mCherry | 23.0 ± 2.4 | - |
| Ppldcp3-eGFP-5 | mEGFP | 8.9 ± 2.8 | ++ |
| Ppldcp3-eGFP-29 | mEGFP | 8.1 ± 1.5 | + |
| Ppldcp3-eGFP-8 | mEGFP | 9.9 ± 3.0 | - |

**Figure S4. Generation and verification of PpLDCP3 translational fusion lines**

(a) – Construction of PpLDCP3-mCh translational fusion in locus.

(b) - Construction of PpLDCP3-mEGFP translational fusion in locus.

(c) – Three lines of the fusion lines (mCherry, mEGFP) were verified for the correct insertion by PCR. In the table in the right, the predicted lengths of amplified regions are shown.

(d) – The percentage of gametophores with sporophytes on the 60$^{th}$ dpi in the selected PpLDCP3 fluorescence lines. The right column contains information about the signal intensity and sporophytes amount in the developing egg cell of the selected lines. The ranking is performed arbitrarily, based on the epifluorescence microscope settings (intensity) needed to detect the signal.
