## Supplemental Figure S5 for "The LOTUS-domain containing protein PpLDCP3 controls germline and dispersal unit formation in the moss *Physcomitrium patens*"

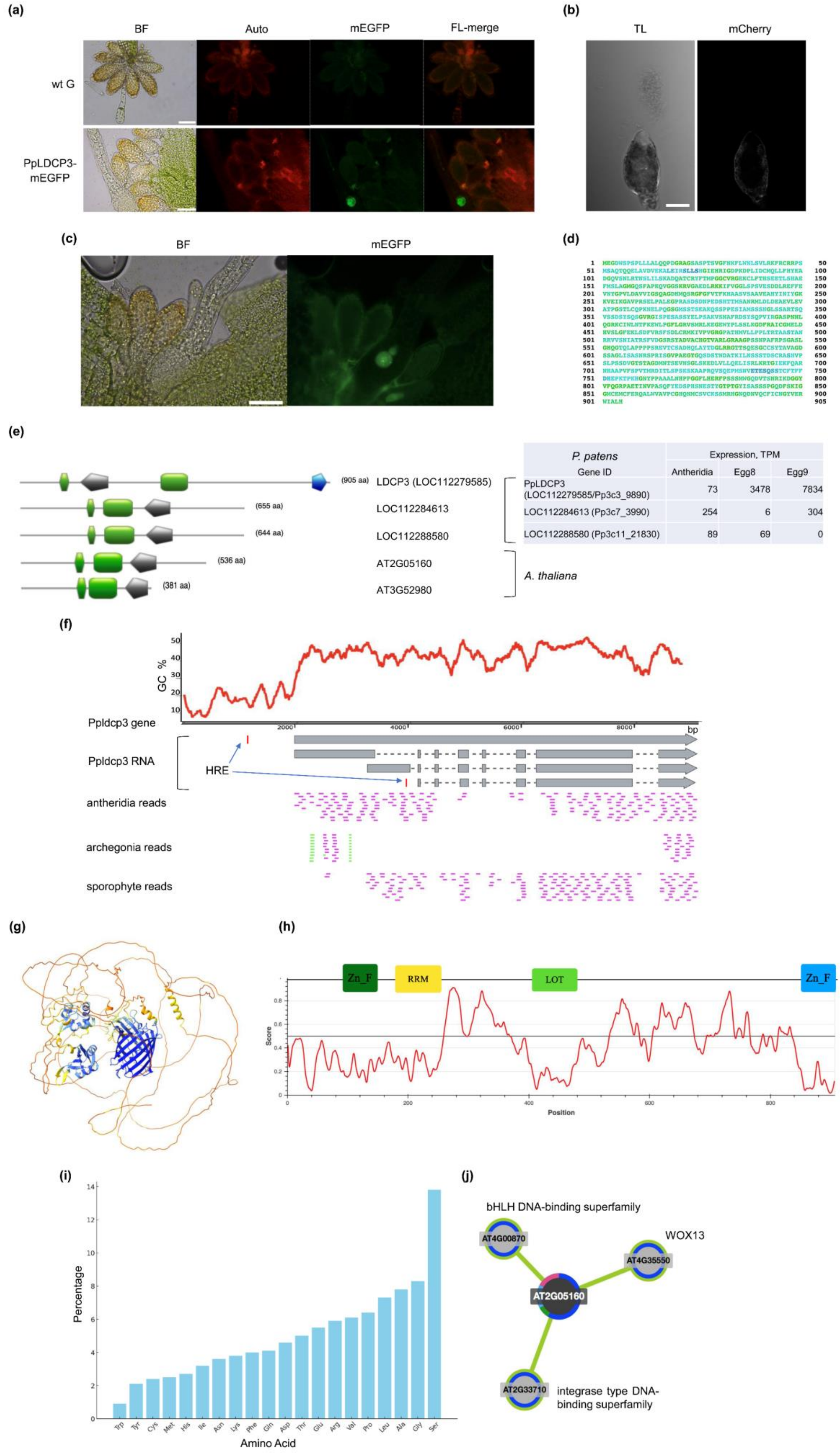
(a)
BF
Auto
mEGFP
FL-merge
wt G
PpLDCP3-mEGFP
(b)
TL
mCherry
(c)
BF
mEGFP
(d)
(e)
(905 aa)
(655 aa)
(644 aa)
(536 aa)
(381 aa)
LDCP3 (LOC112279585)
LOC112284613
LOC112288580
AT2G05160
AT3G52980
A. thaliana
P. patens
Gene ID
Expression, TPM
Antheridia
Egg8
Egg9
PpLDCP3 (LOC112279585/Pp3c3_9890) 73 3478 7834
LOC112284613 (Pp3c7_3990) 254 6 304
LOC112288580 (Pp3c11_21830) 89 69 0
(f)
GC %
Ppldcp3 gene
Ppldcp3 RNA
HRE
antheridia reads
archegonia reads
sporophyte reads
bp
(g)
(h)
Zn_F
RRM
LOT
Zn_F
Score
Position
(i)
Percentage
Amino Acid
Trp
Tyr
Cys
Met
His
Ile
Asn
Lys
Phe
Gln
Asp
Thr
Glu
Arg
Val
Pro
Leu
Ala
Gly
Ser
(j)
bHLH DNA-binding superfamily
AT4G00870
WOX13
AT4G35550
AT2G05160
AT2G33710
integrase type DNA-binding superfamily

**Figure S5. Localization of PpLDCP3 in gametangia and its predicted genetic and biochemical properties, responsible for germline and dispersal unit formation**

(a) – Observation of epifluorescence in wild-type and PpLDCP3-mEGFP line gametangia and adjacent leaves. The scale bar is 50 µm.

(b) – PpLDCPF3-mCh fluorescence in antheridium drops soon after the release of the sperm as determined by confocal microscopy. No specific fluorescence signal was found in the released sperm. The scale bar is 30 µm.

(c) – P-bodies were found at the later stages of archegonia development in the PpLDCP3-mEGFP line when the overall fluorescence signal was observed only in the egg cell and ventral canal cell. The scale bar is 50 µm.

(d) – NucPred verification for PpLDCP3 presence in the nucleus provides support for promiscuous cellular localization of the protein, present both in the nucleus and cytoplasm. NucPred score is 0.53, color code according to NucPred site graphics, with a gradient from blue to red towards a higher probability of the presence in the nuclear compartment.

(e) – *A. thaliana* proteins with domain architecture similar to PpLDCP3. No evident paralogs in the genome of *P. patens* can be found for PpLDCP3. However, proteins with similar domain sets are encoded by 2 genes. The proteins were identified through conserved domain structure search and using the PpLDCP3 LOTUS domain for BLAST NCBI search against *P. patens* and *A. thaliana* proteomes. Domain prediction was done using PROSITE and InterProScan, pictograms correspond to the one in Figure 1 (main text). The longest isoforms of the proteins are represented. The table on the right panel represents the transcriptomics data of the corresponding *P. patens* LDCPs, similar to PpLDCP3 domain architectures. Expression values in *P. patens* gametangia are derived from Sanchez-Vera et al. (2022).

(f) – representation of Ppldcp3 locus (NCBI annotation: LOC112279585; corresponds to Pp3c_9890V3.1; Pp1s74_34V6.1 in other P. patens annotations) with 2000 bp upstream region. The gene is about 6000 bp long, with three transcript variants predicted by NCBI annotation to be expressed from the locus. On top of the diagram, the representation of GC content reveals the regions of high prevalence of AT nucleotides, pointing to the accessibility of the region to RNA polymerases due to low melting temperatures. Vertical red bars indicate the locations of heat-response elements (HRE), corresponding to the consensus sequence "nTTCnnGAAn".

The lower panel shows nucleotide BLAST (https://blast.ncbi.nlm.nih.gov/Blast.cgi) alignment pattern for raw RNA-Seq reads, obtained for antheridia, archegonia, and green sporophyte against Ppldcp3 locus. RNA-Seq data expression values are based on the EVOREPRO database (Julca et al. 2021). Color code for aligned reads corresponds to the BLAST similarity score color scheme.

(g) - AlphaFold prediction of the structure of PpLDCP3-mCh. Color codes reflect confidence in the prediction, generated with the “colour bfactor #1 palette "alphafold”” command in ChimeraX.

(h) – Intrinsic disorder predicted for PpLDCP3 longest isoform, using IUPred prediction server.

(i) – Barplot, visualizing the percentage of amino acids present in the longest isoform of LDCP3 and revealing a strong overrepresentation of the hydroxymethyl group containing serine, the possible site of protein phosphorylation. Calculation and representation by PredictProtein tool (https://predictprotein.org/)

(j) – ThaleMine interaction network for *A. thaliana* AT2G05160.
