## Supplemental Table S1 for "The LOTUS-domain containing protein PpLDCP3 controls germline and dispersal unit formation in the moss *Physcomitrium patens*"

**Table S1. Predicted binding sites for WOX13 and bHLH transcription factors in the promotor region of the Ppldcp3 gene**.

The -2000 bp region from the transcription start site (TSS) of the longest isoform of Ppldcp3 (XM_024514182.1) was considered. The binding sites are following the predicted sites for the *Arabidopsis thaliana* genome.

| Motif sequence | DNA motif consensus | *Arabidopsis thaliana* transcription factor associated with motif | Position of binding site from TSS of Ppldcp3 | Reference |
|---|---|---|---|---|
| TTAATCA | TRATTRR | WOX13 | -103 | (Ikeuchi et al. 2022) |
| CCACTA | CCACBW | WOX13 | -207 | (Ikeuchi et al. 2022) |
| TTAATCA | TRATTRR | WOX13 | -308 | (Ikeuchi et al. 2022) |
| CATATG | CANNTG | WOX13 | -357 | (Ikeuchi et al. 2022) |
| AAAGAAAG | ARAGAVAR | WOX13 | -405 | (Ikeuchi et al. 2022) |
| TTAATCA | TRATTRR | WOX13 | -921 | (Ikeuchi et al. 2022) |
| TAATTGG | TRATTRR | WOX13 | -1101 | (Ikeuchi et al. 2022) |
| TAATTGA | TRATTRR | WOX13 | -1194 | (Ikeuchi et al. 2022) |
| CAAATG | CANNTG | Canonical E-box, bHLH | -1574 | (Toledo-Ortiz et al. 2003) |
| TTAATTA | TRATTRR | WOX13 | -1819 | (Ikeuchi et al. 2022) |
| CATATG | CANNTG | Canonical E-box, bHLH | -1875 | (Toledo-Ortiz et al. 2003) |
