## Supplemental Table S2 for "The LOTUS-domain containing protein PpLDCP3 controls germline and dispersal unit formation in the moss *Physcomitrium patens*"

### Table S1. Primers used in this study

| primer id | primer sequence (5´ -> 3´) | purpose |
|---|---|---|
| AK63 | CGACGTACGTTCGAACAATTGGTTTaaaccagggccatacagagaag | left arm amplification, for Ppldcp3 knock-out |
| AK70 | AGCTAATTACCCTGTTATCCCTAGGgatgtaacaactttggcaattacg | left arm amplification, for Ppldcp3 knock-out |
| AK64 | GCCACGCGTGATATCATGCATGTTaacgtgtgataagttggcgtgc | right arm amplification, for Ppldcp3 knock-out |
| AK65 | AGGCGCGCCCATGGATCGATGTTaaccgcagaactccaacagttg | right arm amplification, for Ppldcp3 knock-out |
| AK118 | agaccctttgatagactgcatgc | Verification Ppldcp3 knock-out |
| AK119 | ttgcagaaattgtgcacgacac | Verification Ppldcp3 knock-out |
| AK120 | tcagagcaatatgtggtatggagc | Verification Ppldcp3 knock-out |
| AK121 | gcaggattcggacgattatcttc | Verification Ppldcp3 knock-out |
| AK72 | ttggaatatcgtactgaaaggcc | Verification Ppldcp3 knock-out |
| oAT43 | AGGGTTTCGCTCATGTGTTG | Verification Ppldcp3 knock-out |
| AL028 | TGTGTGAGTAGTTCCCAGATAAGG | Verification PpLDCP3 translational fusion |
| AK149 | GTATAATAAAGTTGCTacaaagatacatgatccttctaacac | right flank for PpLDCP3 translational fusion |
| AK150 | GAAAGCTGGGTAGGCGCCctgtcaattcgtccttaacac | right flank for PpLDCP3 translational fusion |
| AK151 | GGGGACAACTTTGTATAATAAAGTTGCT | right flank for PpLDCP3 translational fusion |
| C30 | GGGGACCACTTTGTACAAGAAAGCTGGGTA | right flank forPpLDCP3 translational fusion |
| AK146 | AAAAAGCAGGCTGGCGCCctatcctcctgctgctgctcttaatc | left flank for PpLDCP3 translational fusion |
| AK147 | TTGTATACAAAGTTGTatgcagggcaatccatctctcc | left flank for PpLDCP3 translational fusion |
| C29 | GGGGACAAGTTTGTACAAAAAAGCAGGCT | left flank for PpLDCP3 translational fusion |
| AK148 | GGGGACAACTTTTGTATACAAAGTTGT | left flank for PpLDCP3 translational fusion |
